# Complement component 4 genes contribute sex-specific vulnerability in diverse illnesses

**DOI:** 10.1101/761718

**Authors:** Nolan Kamitaki, Aswin Sekar, Robert E. Handsaker, Heather de Rivera, Katherine Tooley, David L. Morris, Kimberly E. Taylor, Christopher W. Whelan, Philip Tombleson, Loes M. Olde Loohuis, Schizophrenia Working Group of the Psychiatric Genomics Consortium, Michael Boehnke, Robert P. Kimberly, Kenneth M. Kaufman, John B. Harley, Carl D. Langefeld, Christine E. Seidman, Michele T. Pato, Carlos N. Pato, Roel A. Ophoff, Robert R. Graham, Lindsey A. Criswell, Timothy J. Vyse, Steven A. McCarroll

**Affiliations:** Department of Genetics, Harvard Medical School, Boston, Massachusetts 02115, USA; Stanley Center for Psychiatric Research, Broad Institute of MIT and Harvard, Cambridge, Massachusetts 02142, USA; Department of Medical and Molecular Genetics, King’s College London, London WC2R 2LS, UK; Rosalind Russell / Ephraim P Engleman Rheumatology Research Center, Division of Rheumatology, UCSF School of Medicine, San Francisco, California 94143, USA; Department of Human Genetics, David Geffen School of Medicine, University of California, Los Angeles, California 90095, USA; Center for Neurobehavioral Genetics, Semel Institute for Neuroscience and Human Behavior, University of California, Los Angeles, California 90095, USA; A full list of collaborators is in Supplementary Information; Department of Biostatistics and Center for Statistical Genetics, University of Michigan, Ann Arbor, Michigan 48109, USA; Division of Clinical Immunology and Rheumatology, University of Alabama at Birmingham, Birmingham, Alabama 35294, USA; Center for Autoimmune Genomics and Etiology (CAGE), Department of Pediatrics, Cincinnati Children’s Medical Center & University of Cincinnati and the US Department of Veterans Affairs Medical Center, Cincinnati, Ohio, USA; Department of Biostatistical Sciences, Wake Forest School of Medicine, Winston-Salem, North Carolina 27101, USA; Howard Hughes Medical Institute, Chevy Chase, Maryland 20815, USA; Cardiovascular Division, Brigham and Women’s Hospital, Boston, Massachusetts 02115, USA; SUNY Downstate Medical Center, Brooklyn, New York 11203, USA; Human Genetics, Genentech Inc, South San Francisco, California 94080, USA

## Abstract

Many common illnesses differentially affect men and women for unknown reasons. The autoimmune diseases lupus and Sjögren’s syndrome affect nine times more women than men^1,2^, whereas schizophrenia affects men more frequently and severely^3–5^. All three illnesses have their strongest common-genetic associations in the Major Histocompatibility Complex (MHC) locus, an association that in lupus and Sjögren’s syndrome has long been thought to arise from *HLA* alleles^6–13^. Here we show that the complement component 4 (*C4*) genes in the MHC locus, recently found to increase risk for schizophrenia^14^, generate 7-fold variation in risk for lupus (95% CI: 5.88-8.61; *p* < 10^−117^ in total) and 16-fold variation in risk for Sjögren’s syndrome (95% CI: 8.59-30.89; *p* < 10^−23^ in total), with *C4A* protecting more strongly than *C4B* in both illnesses. The same alleles that increase risk for schizophrenia, greatly reduced risk for lupus and Sjögren’s syndrome. In all three illnesses, *C4* alleles acted more strongly in men than in women: common combinations of *C4A* and *C4B* generated 14-fold variation in risk for lupus and 31-fold variation in risk for Sjögren’s syndrome in men (vs. 6-fold and 15-fold among women respectively) and affected schizophrenia risk about twice as strongly in men as in women. At a protein level, both C4 and its effector (C3) were present at greater levels in men than women in cerebrospinal fluid (*p* < 10^−5^ for both C4 and C3) and plasma among adults ages 20-50^15–17^, corresponding to the ages of differential disease vulnerability. Sex differences in complement protein levels may help explain the larger effects of *C4* alleles in men, women’s greater risk of SLE and Sjögren’s, and men’s greater vulnerability in schizophrenia. These results nominate the complement system as a source of sexual dimorphism in vulnerability to diverse illnesses.

---

Systemic lupus erythematosus (SLE, or “lupus”) is a systemic autoimmune disease of unknown cause. Risk of SLE is heritable (66%^18^), though SLE may have environmental triggers, as its onset often follows events that damage cells, such as infections and severe sunburns^19^. Most SLE patients produce autoantibodies against nucleic acid complexes, including ribonucleoproteins and DNA^20^.

In genetic studies, SLE associates most strongly with variation across the major histocompatibility complex (MHC) locus^6,7,21^. However, conclusive attribution of this association to specific genes and alleles has been difficult; the identities of the most likely genic and allelic culprits have been frequently revised as genetic studies have grown in size^8–11^. In several other autoimmune diseases, including type 1 diabetes, celiac disease, and rheumatoid arthritis, strong effects of the MHC locus arise from *HLA* alleles that cause the peptide binding groove of HLA proteins to present a disease-critical autoantigen^22–24^. In SLE, by contrast, MHC alleles associate broadly with the presence of diverse autoantibodies^25^.

The complement component 4 (*C4A* and *C4B*) genes are also present in the MHC locus, between the class I and class II *HLA* genes. Classical complement proteins help eliminate debris from dead and damaged cells, attenuating the exposure of diverse intracellular proteins to the adaptive immune system. *C4A* and *C4B* commonly vary in genomic copy number^26–28^ and encode complement proteins with distinct affinities for molecular targets^29,30^. SLE frequently presents with hypocomplementemia that worsens during flares, possibly reflecting increased active consumption of complement^31^. Rare cases of severe, early-onset SLE can involve complete deficiency of a complement component (C4, C2, or C1Q)^32,33^, and one of the strongest common-variant associations in SLE maps to *ITGAM*, which encodes a receptor for C3, the downstream effector of C4^21,34^. Though total *C4* gene copy number associates with SLE risk^35–37^, this association is thought to arise from linkage disequilibrium (LD) with nearby *HLA* alleles^38^, which have been the focus of fine-mapping analyses^6–11,21^.

The complex genetic variation at *C4* – arising from many alleles with different numbers of *C4A* and *C4B* genes – has been challenging to analyze in large cohorts. A recently feasible approach to this problem is based on imputation: people share long haplotypes with the same combinations of SNP and *C4* alleles, such that *C4A* and *C4B* gene copy numbers can be imputed from SNP data^14^. To analyze *C4* in large cohorts, we developed a way to identify *C4* alleles from whole-genome sequence (WGS) data (**Fig. 1**), then analyzed WGS data from 1,265 individuals (from the Genomic Psychiatry Cohort^39,40^) to create a large multi-ancestry panel of 2,530 reference haplotypes of MHC SNPs and *C4* alleles (**Extended Data Fig. 1**) – ten times more than in earlier work^14^. We then analyzed SNP data from the largest SLE genetic association study^7^ (ImmunoChip 6,748 SLE cases and 11,516 controls of European ancestry) (**Extended Data Fig. 2**), imputing *C4* alleles to estimate the SLE risk associated with common combinations of *C4A* and *C4B* gene copy numbers (**Fig. 2a**).

**Figure 1.**
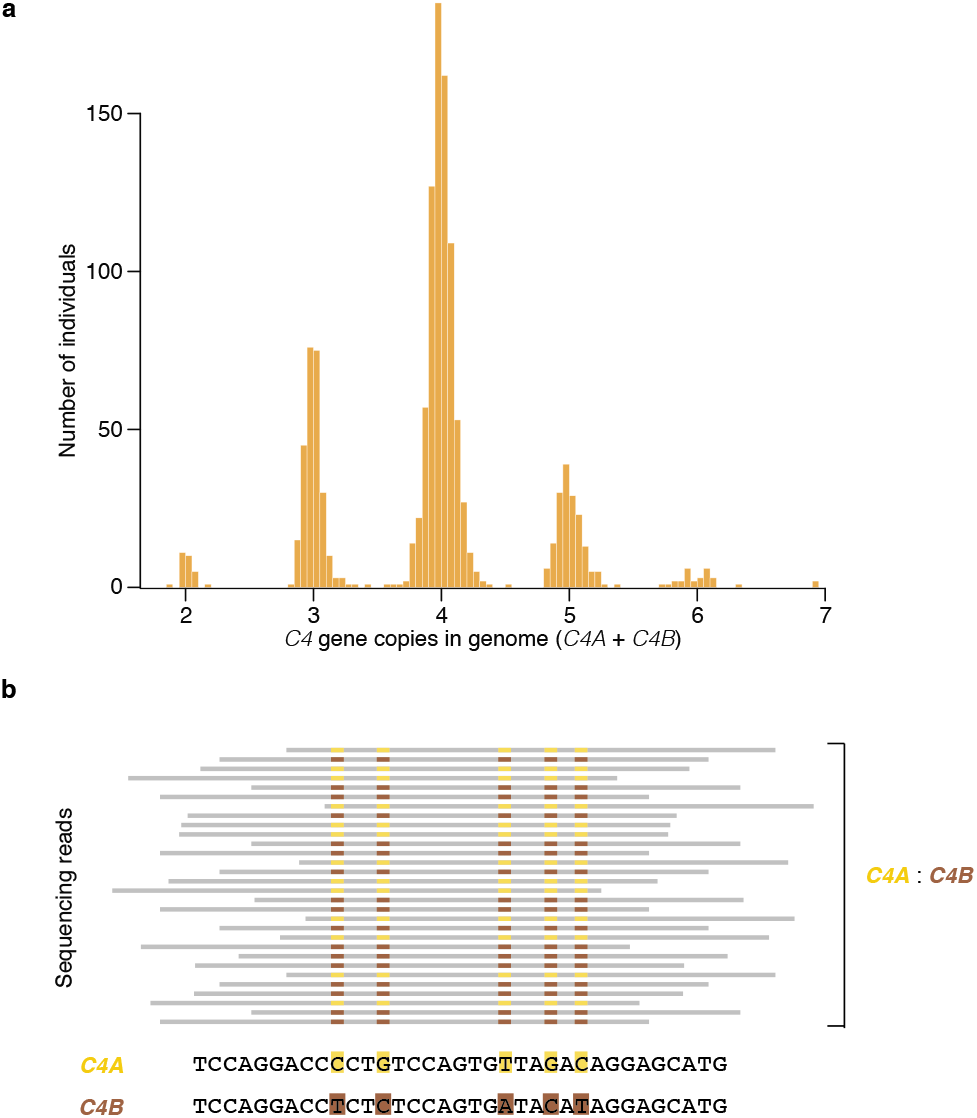
Analysis of *C4* gene variation by whole-genome sequencing. (a) Distributions (across 1,265 individuals) of total *C4* gene copy number (*C4A* + *C4B*), as measured from read depth of coverage across the *C4* locus, in whole-genome sequencing data. (b) The relative numbers of reads that overlap sequences specific to *C4A* or *C4B* (together with the total C4 gene copy number, **a**) are used to infer the underlying copy numbers of the *C4A* and *C4B* genes. For example, in an individual with four *C4* genes, the presence of equal numbers of reads specific to *C4A* or *C4B* suggests the presence of two copies each of *C4A* and *C4B*. Precise statistical approaches (including inference of probabilistic dosages), and further approaches for phasing *C4* allelic states with nearby SNPs to create reference haplotypes, are described in **Methods**.

**Figure 2.**
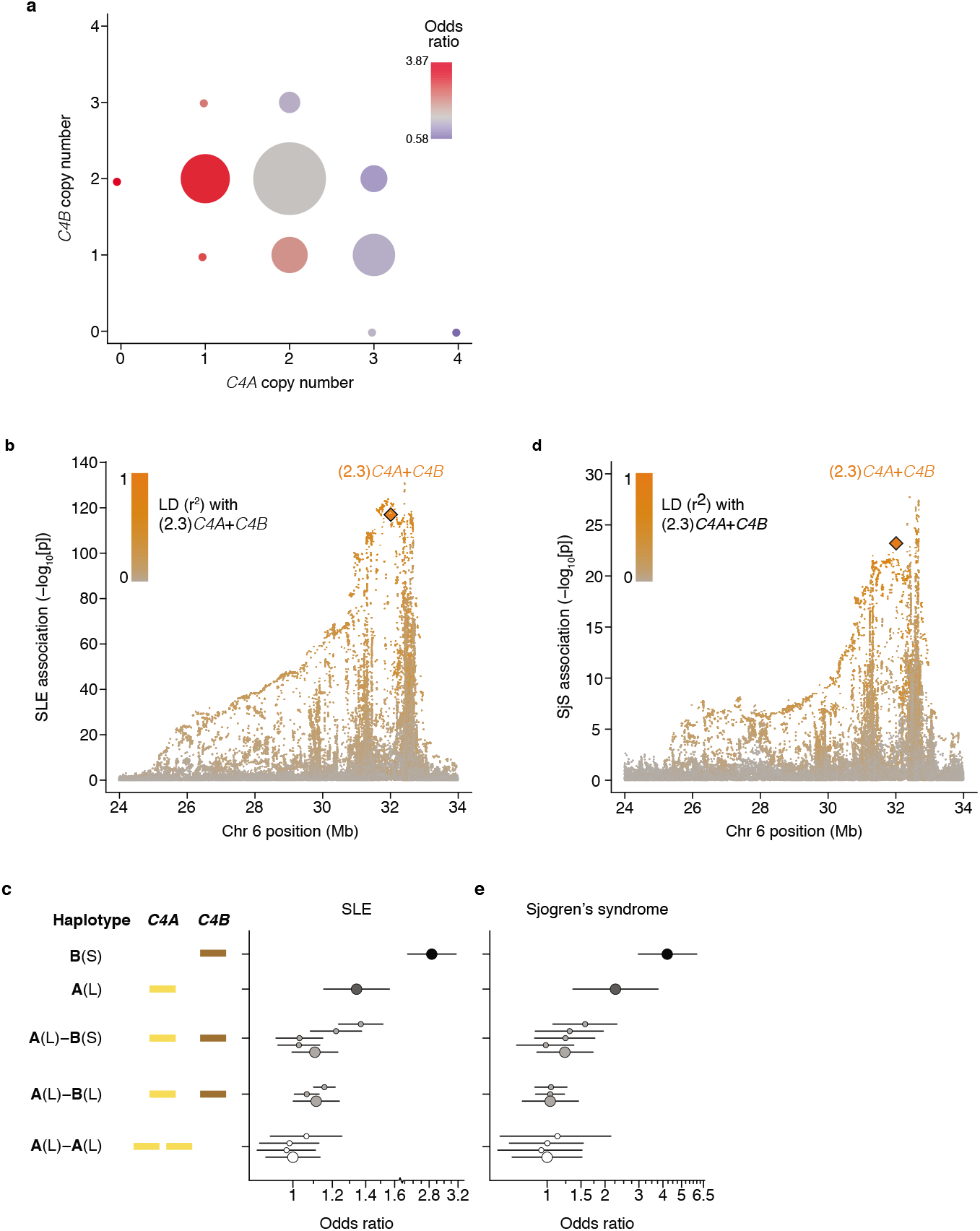
Association of SLE with *C4* alleles. (a) Levels of SLE risk associated with 11 common combinations of *C4A* and *C4B* gene copy number. The color of each circle reflects the level of SLE risk (odds ratio) associated with a specific combination of *C4A* and *C4B* gene copy numbers relative to the most common combination (two copies of *C4A* and two copies of *C4B*) in gray. The area of each circle is proportional to the number of individuals with that number of *C4A* and *C4B* genes. Paths from left to right on the plot reflect the effect of increasing *C4A* gene copy number; paths from bottom to top reflect the effect of increasing *C4B* gene copy number; and diagonal paths from upper left to lower right reflect the effect of exchanging *C4B* for *C4A* copies. Data are from analysis of 6,748 SLE cases and 11,516 controls of European ancestry. The odds ratios are reported with confidence intervals in **Extended Data Fig. 3**. (b) Association of SLE with genetic markers (SNPs and imputed *HLA* alleles) across the extended MHC locus within the European-ancestry cohort. Orange diamond: an initial estimate of *C4*-related genetic risk, calculated as a weighted sum of the number of *C4A* and *C4B* gene copies: (2.3)*C4A*+*C4B*, with the weights derived from the relative coefficients estimated from logistic regression of SLE risk vs. *C4A* and *C4B* gene dosages. This risk score is imputed with an accuracy (*r*^2^) of 0.77. Points representing all other genetic variants in the MHC locus are shaded orange according to their level of linkage disequilibrium–based correlation to this *C4*-derived risk score. (c) SLE risk associated with common combinations of *C4* structural allele and MHC SNP haplotype. For each *C4* locus structure, separate odds ratios are reported for each “haplogroup,” i.e., the MHC SNP haplotype background on which the *C4* structure segregates. Error bars represent 95% confidence intervals around the effect size estimate for each sex. (d) As in (b), but with a cohort of 673 Sjögren’s Syndrome (SjS) cases and 1,153 controls of European ancestry. The orange diamond is also an estimate of *C4*-related genetic risk calculated as a weighted sum of *C4A* and *C4B* gene copies estimated from a logistic regression of SjS risk: (2.3)*C4A*+*C4B*. (e) As in (c), but with the SjS cohort from (d). Error bars represent 95% confidence intervals around the effect size estimate for each sex.

Groups with the eleven most common combinations of *C4A* and *C4B* gene copy number exhibited 7-fold variation in their risk of SLE (**Fig. 2a** and **Extended Data Fig. 3**). The relationship between SLE vulnerability and *C4* gene copy number exhibited consistent, logical patterns across the 11 genotype groups. For each *C4B* copy number, greater *C4A* copy number associated with reduced SLE risk (**Fig. 2a, Extended Data Fig. 3**). For each *C4A* copy number, greater *C4B* copy number associated with more modestly reduced risk (**Fig. 2a**). Logistic-regression analysis estimated that the protection afforded by each copy of *C4A* (OR: 0.54; 95% CI: [0.51, 0.57]) was equivalent to that of 2.3 copies of *C4B* (OR: 0.77; 95% CI: [0.71,0.82]). We calculated an initial *C4*-derived risk score as 2.3 times the number of *C4A* genes, plus the number of *C4B* genes, in an individual’s genome. Despite clear limitations of this risk score – it is imperfectly imputed from flanking SNP haplotypes (*r*^2^ = 0.77, **Extended Data Table 1**) and only approximates *C4*-derived risk by using a simple, linear model (to avoid over-fitting the genetic data) – SNPs across the MHC locus tended to associate with SLE in proportion to their level of LD with this risk score (**Fig. 2b**).

Combinations of many different *C4* alleles generate the observed variation in *C4A* and *C4B* gene copy number; particular *C4A* and *C4B* gene copy numbers have also arisen recurrently on multiple SNP haplotypes^14^ (**Extended Data Fig. 1**). Analysis of SLE risk in relation to each of these alleles and haplotypes reinforced the conclusion that *C4A* contributes strong protection, and *C4B* more modest protection, from SLE, and that *C4* genes (rather than nearby variants) are the principal drivers of this variation in risk levels (**Fig. 2c**).

These results prompted us to consider whether other autoimmune disorders with similar patterns of genetic association at the MHC locus might also be driven in part by *C4* variation. Sjögren’s syndrome (SjS) is a heritable (54%^41^) systemic autoimmune disorder of exocrine glands, characterized primarily by dry eyes and mouth with other systemic effects. At a protein level, SjS is (like SLE) characterized by diverse autoantibodies^42^, including antinuclear antibodies targeting ribonucleoproteins^43^, and hypocomplementemia^44,45^. The largest source of common genetic risk for SjS lies in the MHC locus^46^, with associations to the same haplotype(s) as in SLE^12,13^ and with heterogeneous *HLA* associations in different ancestries^47^. We imputed *C4* alleles into existing SNP data from a European-ancestry SjS case-control cohort (673 cases and 1153 controls). As in SLE, logistic-regression analyses found both *C4A* copy number (OR: 0.41; 95% CI: [0.34, 0.49]) and *C4B* copy number (OR: 0.67; 95% CI: [0.53, 0.86]) to be protective against SjS. The risk-equivalent ratio of *C4B* to *C4A* gene copies was similar in SjS and SLE (about 2.3 to 1); also, as with SLE, nearby SNPs associated with SjS in proportion to their LD with a *C4*-derived risk score ((2.3)*C4A*+*C4B*) (**Fig. 2d**). The distribution of SjS risk across the individual *C4* alleles and haplotypes revealed a pattern that (as in SLE) supported greater protective effect from *C4A* than *C4B*, and little effect of flanking SNP haplotypes (**Fig. 2e**).

The association of SLE and SjS with *C4* gene copy number has long been attributed to the *HLA-DRB1*03:01* allele. In European populations, *DRB1*03:01* is in strong LD (*r*^2^ = 0.71) with the common *C4*-**B**(S) allele, which lacks any *C4A* gene and is the highest-risk *C4* allele in our analysis (**Fig. 2c**); many MHC SNPs associated with SLE and SjS in proportion to their LD correlations with both *C4* and DRB1*03:01 (**Extended Data Fig. 4a, b**). Cohorts with other ancestries can have recombinant haplotypes that disambiguate the contributions of alleles that are in LD in Europeans. Among African Americans, we found that common *C4* alleles exhibited far less LD with *HLA* alleles; in particular, the LD between *C4*-B(S) and *DRB1*03:01* was low (*r*^2^ = 0.10) (**Extended Data Table 2**). Thus, genetic data from an African American SLE cohort (1,494 cases, 5,908 controls) made it possible to distinguish between these potential genetic effects. Joint association analysis of C4A, *C4B*, and *DRB1*0301* implicated *C4A* (*p* < 10^−14^) and *C4B* (*p* < 10^−5^) but not *DRB1*0301* (*p* = 0.29) (**Extended Data Table 3**). Each *C4* allele associated with effect sizes of similar magnitude on SLE risk in Europeans and African Americans (**Fig. 3a**). An analysis specifically of combinations of *C4*-**B**(S) and *DRB1*\*03:01 allele dosages in African Americans showed that *C4*-**B**(S) alleles consistently increased SLE risk regardless of *DRB1*\*03:01 status, whereas *DRB1*\*03:01 had no consistent effect when controlling for *C4*-**B**(S) (**Fig. 3b**). Although *C4* alleles had less LD with nearby variants on African American than on European haplotypes, SNPs associated with SLE in proportion to LD correlations with *C4* in African Americans as well (**Extended Data Fig. 4c**).

**Figure 3.**
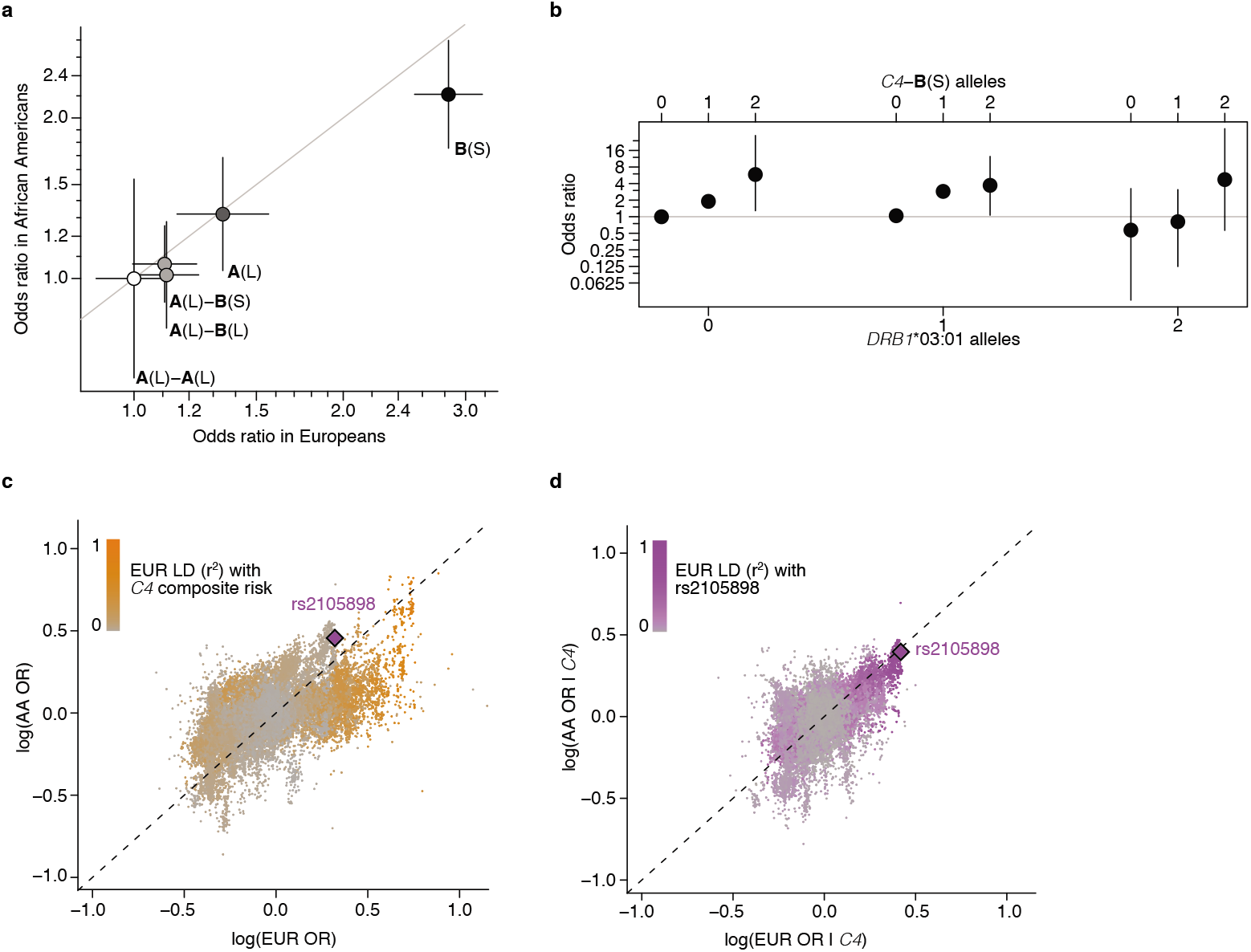
*C4* and trans-ancestral analysis of the MHC association signal in SLE. (a) Common *C4* alleles exhibit similar strengths of association (odds ratios) in European ancestry and African American (1,494 SLE cases; 5,908 controls) cohorts. Error bars represent 95% confidence intervals around the effect size estimate for each sex. (b) Analysis of SLE risk across combinations of *C4*-**B**(S) and *DRB1*\*03:01 genotypes in an African American SLE case–control cohort, in which the two alleles exhibit very little LD (*r*^2^ = 0.10). On each *DRB1*\*03:01 genotype background, additional *C4*-**B**(S) alleles increase risk (ie. within each grouping). Whereas on each C4-**B**(S) background, *DRB1*\*03:01 alleles have no appreciable relationship with risk (ie. every n^th^ point from each group). Error bars represent 95% confidence intervals around the effect size estimate for each combination of *C4*-**B**(S) and *DRB1*\*03:01. (c) Trans-ancestry comparison of the association of genetic markers with SLE (unconditioned log-odds ratios) among European-ancestry (x-axis) and African American (y-axis) research participants. LD with *C4*-derived risk in European-ancestry individuals (indicated by orange shading) contributes to the apparent discordance of association patterns between populations. A lead SNP identified below, rs2105898 (purple), is among the strongest signals in the African American cohort; among Europeans, though, its association is initially much less remarkable than that of other SNPs that are in strong LD with *C4*. (d) In analyses controlling for *C4*-derived risk, analyses of European ancestry and African American cohorts both identified a small haplotype (tagged by rs2105898) harboring a genetic signal independent of *C4*. Several SNPs that form a short haplotype common to both ancestry groups are among the top associations in both cohorts. Further analyses of this haplotype are described in **Supplementary Note**. Many SNP associations that appear specific to the European-ancestry cohort have European-ancestry LD with rs2105898 (purple shading) in excess of LD with the same haplotype in the African American cohort (**Extended Data Fig. 8**).

We next sought to find other potential contributions of the MHC locus to SLE risk by accounting for contributions from *C4*. SNPs across the MHC locus display very different associations with SLE in Europeans and African Americans^7,11^, though the SNPs with European-specific associations tend to have strong LD to *C4* in Europeans (**Fig. 3c**). To control for *C4* genotypes, many of which exhibit strong LD across the MHC locus in Europeans (**Extended Data Fig. 1**), we adjusted the association data for *C4*-derived risk using a more-complete *C4*-derived risk score derived from the genotype-group risk measurements in **Fig. 2a**. Once adjusted for C4 effects, the residual association signals in the two populations became strongly correlated (**Fig. 3d**). Both populations also pointed to the same small haplotype of two variants as the most likely driver of an additional genetic effect independent of *C4* (**Fig. 3d** and **Supplementary Note**). The two variants defining this short haplotype reside within the XL9 regulatory region^48,49^, a well-studied region of open chromatin that contains abundant chromatin marks characteristic of active enhancers and transcription factor binding sites (**Supplementary Note**). One of these variants, rs2105898, disrupts a binding site for ZNF143^50^, which anchors interactions of distal enhancers with gene promoters^51^ (**Supplementary Note**). Data from the GTEx Consortium ^52^ (v7) included 227 instances (gene/tissue pairs) in which this haplotype associated with elevated (*HLA-DRB1, -DRB5, -DQA1*, and *-DQB1*) or reduced (*HLA-DRB6, -DQA2*, and *-DQB2*) expression of an *HLA* class II gene with at least nominal (*p* < 10^−4^) significance. Some of the strongest associations at each gene (*p* < 10^−8^ to 10^−76^) were in whole blood, but expression QTLs elsewhere can also reflect the presence of blood and immune cells within those tissues.^53^ (Although eQTL analyses of *HLA* genes may be affected by read-alignment artifacts in these genes’ hyperpolymorphic domains, most such observed signals are robust after adjusting for individual *HLA* alleles.^54^)

The haplotype with elevated expression of *HLA-DRB1, -DRB5, -DQA1*, and *-DQB1* (allele frequency 0.20 among Europeans, 0.22 among African Americans) associated with increased SLE risk (odds ratio) of 1.52 (95% CI: 1.44-1.61;*p* < 10^−48^) in Europeans and 1.49 (95% CI: 1.35-1.63; *p* < 10^−16^) in African Americans in analyses adjusting for *C4* effects. The risk haplotype was in strong LD with *DRB1*\*15:01 in Europeans and *DRB1*\*15:03 in African Americans, which may explain earlier findings of population-specific associations with *DRB1*\*15:01 in Europeans and *DRB1*\*15:03 in African Americans^7,11^. The risk haplotype tagged by rs2105898 tended to be on low-risk *C4* haplotypes in Europeans, a relationship that may have made both genetic influences harder to recognize in earlier work; controlling for either rs2105898 or *C4* (**Extended Data Fig. 5a**) greatly increased the association of SLE with the other genetic influence (**Extended Data Table 3**). Controlling for the simpler (2.3)*C4A*+*C4B* model in SNP associations with SjS (as precision of estimates of individual alleles were low due to sample size) also pointed strongly to the same haplotype, with the same allele of rs2105898 associating in the same direction but larger effect (OR: 1.96; 95% CI: 1.64-2.34) as compared to SLE (**Extended Data Fig. 5b**).

Alleles at *C4* that increase dosage of *C4A*, and to a lesser extent *C4B*, appear to protect strongly against SLE and SjS (**Fig. 2a-c**); by contrast, alleles that increase expression of *C4A* in the brain are more common among individuals with schizophrenia^6^. These same illnesses exhibit striking, and opposite, sex differences: SLE and SjS are nine times more common among women of childbearing age than among men of a similar age^1,2^, whereas in schizophrenia, women exhibit less severe symptoms, more frequent remission of symptoms, lower relapse rates, and lower overall incidence ^3–5^. Hence, we sought to evaluate the possibility that the effects of *C4* alleles on the risk of each disease might also differ between men and women.

Analysis indicated that the effects of *C4* alleles in both lupus and schizophrenia were stronger in men. When a sex-by-*C4* interaction term was included in association analyses, this term was significant for both SLE (*p* < 0.01) and schizophrenia (*p* < 0.01), indicating larger *C4* effects in men for both disorders. (Analysis of SjS had limited power due to the small number of men affected by SjS – 60 of the 673 cases in the cohort – but pointed to the same direction of effect at *p* = 0.07). For both SLE and schizophrenia, the individual *C4* alleles consistently associated with stronger effects in men than women (**Fig. 4a, b**). These relationships explained previously reported sex biases ^55^ in SNP associations across the MHC locus (**Fig. 4c-e**).

**Figure 4.**
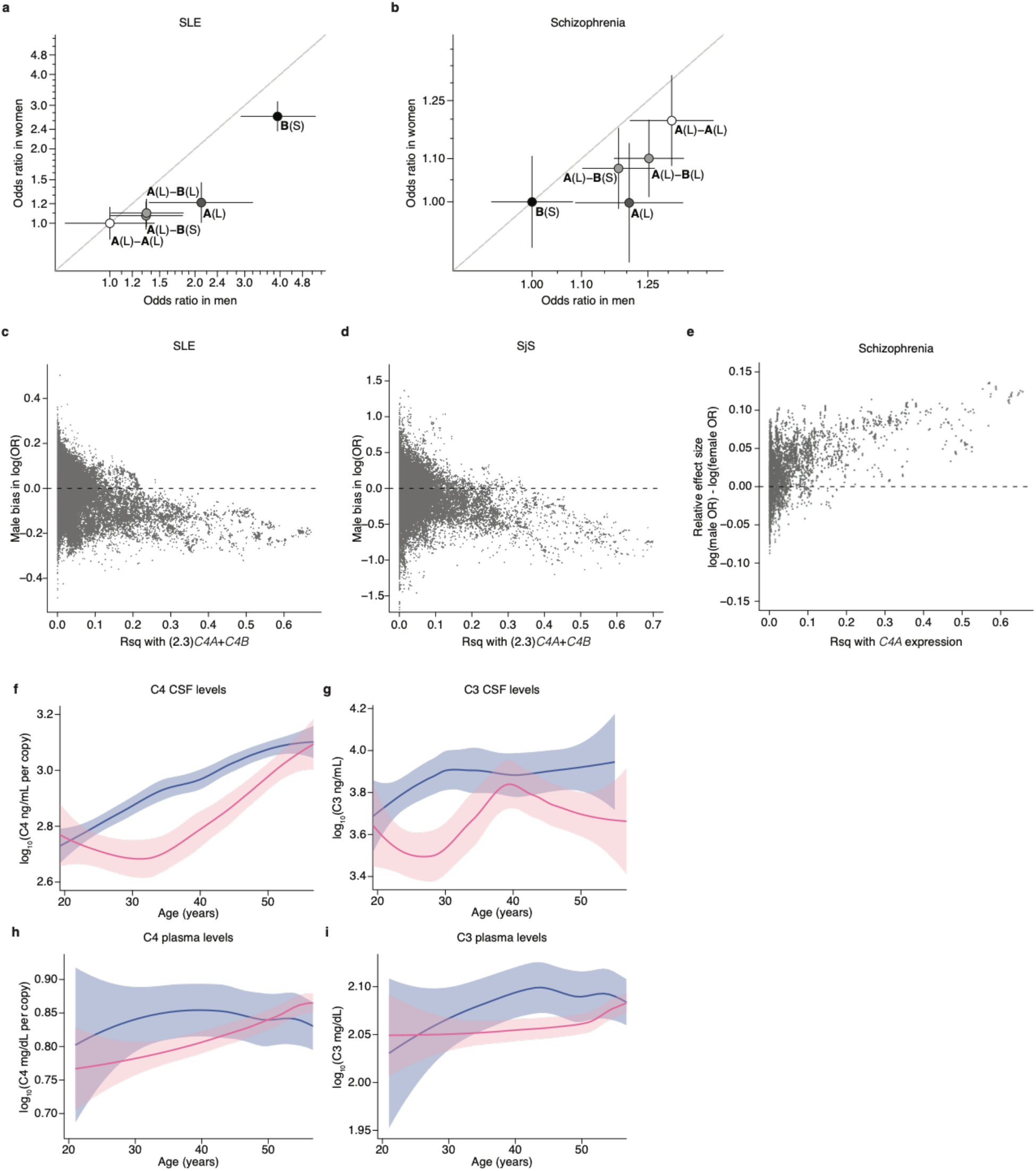
Sex differences in the magnitude of *C4* genetic effects and complement protein concentrations. (a) SLE risk (odds ratios) associated with the four most common *C4* alleles in men (x-axis) and women (y-axis) among 6,748 affected and 11,516 unaffected individuals of European ancestry. For each sex, the lowest-risk allele (*C4*-**A**(L)-**A**(L)) is used as a reference (odds ratio of 1.0). Shading of each allele reflects the relative level of SLE risk conferred by *C4A* and *C4B* copy numbers as in **Fig. 2c**. Error bars represent 95% confidence intervals around the effect size estimate for each sex. (b) Schizophrenia risk (odds ratios) associated with the four most common *C4* alleles in men (x-axis) and women (y-axis) among 28,799 affected and 35,986 unaffected individuals of European ancestry, aggregated by the Psychiatric Genomics Consortium^62^. For each sex, the lowest-risk allele (*C4*-**B**(S)) is used as a reference (odds ratio of 1.0). For visual comparison with **Fig. 4a**, shading of each allele reflects the relative level of SLE risk. Error bars represent 95% confidence intervals around the effect size estimate for each sex. (c) Relationship between male bias in SLE risk (difference between male and female log–odds ratios) and LD with *C4* risk for common (minor allele frequency [MAF] > 0.1) genetic markers across the extended MHC region. For each SNP, the allele for which sex risk bias is plotted is the allele that is positively correlated (via LD) with *C4*-derived risk score. (d) Relationship between male bias in SjS risk (log-odds ratios) and LD with *C4* risk for common (minor allele frequency [MAF] > 0.1) genetic markers across the extended MHC region. For each SNP, the allele for which sex risk bias is plotted is the allele that is positively correlated (via LD) with *C4*-derived risk score. (e) Relationship of male bias in schizophrenia risk (log–odds ratios) and LD to *C4A* expression for common (MAF > 0.1) genetic markers across the extended MHC region. For each SNP, the allele for which sex risk bias is plotted is the allele that is positively correlated (via LD) with imputed *C4A* expression. (f) Concentrations of C4 protein in cerebrospinal fluid sampled from 340 adult men (blue) and 167 adult women (pink) as a function of age with local polynomial regression (LOESS) smoothing. Concentrations are normalized to the number of *C4* gene copies in an individual’s genome (a strong independent source of variance, **Extended Data Figure 7a**) and shown on a log_10_ scale. Shaded regions represent 95% confidence intervals derived during LOESS smoothing. (g) Levels of C3 protein in cerebrospinal fluid from 179 adult men and 125 adult women as a function of age. Concentrations are shown on a log_10_ scale. Shaded regions represent 95% confidence intervals derived during LOESS smoothing. (h) Levels of C4 protein in blood plasma from 182 adult men and 1662 adult women as a function of age. As in (f), concentrations are normalized to C4 gene copy number (**Extended Data Fig. 7b**) and shown on a log_10_ scale. Shaded regions represent 95% confidence intervals derived during LOESS smoothing. (i) Levels of C3 protein in blood plasma as a function of age from the same individuals in **Fig. 4h**. Concentrations are shown on a log_10_ scale. Shaded regions represent 95% confidence intervals derived during LOESS smoothing.

The stronger effects of *C4* alleles on male relative to female risk could arise from sex differences in *C4* RNA expression, C4 protein levels, or downstream responses to C4. Analysis of RNA expression in 45 tissues, using data from GTEx^52^, identified no sex differences in *C4* RNA expression. We then analyzed C4 protein in cerebrospinal fluid (CSF) from two panels of adult research participants (*n* = 589 total) in whom we had also measured *C4* gene copy number by direct genotyping or imputation. CSF C4 protein levels correlated strongly with *C4* gene copy number (*p* < 10^−10^, **Extended Data Fig. 6a**), so we normalized C4 protein measurements to the number of *C4* gene copies. CSF from adult men contained on average 27% more C4 protein per *C4* gene copy than CSF from women (meta-analysis *p* = 9.9 × 10^−6^, **Fig. 4f**). C4 acts by activating the complement component 3 (C3) protein, promoting C3 deposition onto targets in tissues. CSF levels of C3 protein were also on average 42% higher among men than women (meta-analysis *p* = 7.5 × 10^−7^, **Fig. 4g**).

The elevated concentrations of C3 and C4 proteins in CSF of men parallel earlier findings that, in plasma, C3 and C4 are also present at higher levels in men than women^15–17^. The large sample size (n > 50,000) of the plasma studies allows sex differences to be further analyzed as a function of developmental age. Both men and women undergo age-dependent elevation of C4 and C3 levels in plasma, but this occurs early in adulthood (age 20–30) in men and closer to menopause (age 40–50) in women, with the result that male– female differences in complement protein levels are observed primarily during the reproductive years (ages 20–50). We replicated these findings using measurements of C3 and (gene copy number-corrected) (**Extended Data Fig. 6b**) C4 protein in plasma from adults, finding (as in the earlier plasma studies and in CSF) that these differences are most pronounced during the reproductively active years of adulthood (ages 20-50) (**Fig. 4h, i**). We also observed that SjS patients have lower C4 serum levels than controls (*p* < 1×10^−20^, **Extended Data Fig. 6c**) even after correcting for *C4* gene copy number (*p* < 1×10^−8^, **Extended Data Fig. 6d**), suggesting that hypocomplementemia in SjS is not simply due to *C4* genetics but also reflects disease effects on ambient complement levels, for example due to complement consumption. The ages of pronounced sex difference in complement levels corresponded to the ages at which men and women differ in disease incidence: in schizophrenia, men outnumber women among cases incident in early adulthood, but not among cases incident after age 40^4,56^; in SLE, women greatly outnumber men among cases incident during the child-bearing years, but not among cases incident after age 50 or during childhood^57^; in SjS, the large relative vulnerability of women declines in magnitude after age 50 ^58,59^.

Our results indicate that the MHC locus shapes vulnerability in lupus and SjS – two of the three most common rheumatic autoimmune diseases – in a very different way than in type I diabetes, rheumatoid arthritis, and celiac disease. In those diseases, precise interactions between specific *HLA* alleles and specific autoantigens determine risk ^22–24^. In SLE and SjS, however, the genetic variation implicated here points instead to the continuous, chronic interaction of the immune system with very many potential autoantigens. Because complement facilitates the rapid clearance of debris from dead and injured cells, elevated levels of C4 protein likely attenuate interactions between the adaptive immune system and ribonuclear self-antigens at sites of cell injury, pre-empting the development of autoimmunity. The additional *C4*-independent genetic risk effect described here (associated with rs2105898) may also affect autoimmunity broadly, rather than antigen-specifically, by regulating expression of many HLA class II genes (including *DRB1, DQA1*, and *DQB1*). Mouse models of SLE indicate that once tolerance is broken for one self-antigen, autoreactive germinal centers generate B cells targeting other self-antigens^60^; such “epitope spreading” could lead to autoreactivity against many related autoantigens, regardless of which antigen(s) are involved in the earliest interactions with immune cells. Our genetic findings address the development of SLE and SjS rather than complications that arise in any specific organ. A few percent of SLE patients develop neurological complications that can include psychosis^61^; though psychosis is also a symptom of schizophrenia, neurological complications of SLE do not resemble schizophrenia more broadly, and likely have a different etiology.

The same *C4* alleles that increase vulnerability to schizophrenia appeared to protect strongly against SLE and SjS. This pleiotropy will be important to consider in efforts to engage the complement system therapeutically. The complement system contributed to these pleiotropic effects more strongly in men than in women. Moreover, though the allelic series at *C4* allowed human genetics to establish dose-risk relationships for *C4*, sexual dimorphism in the complement system also extended to complement component 3 (C3). Why and how biology has come to create this sexual dimorphism in the complement system in humans presents interesting questions for immune and evolutionary biology.

**Extended Data Figure 1.**
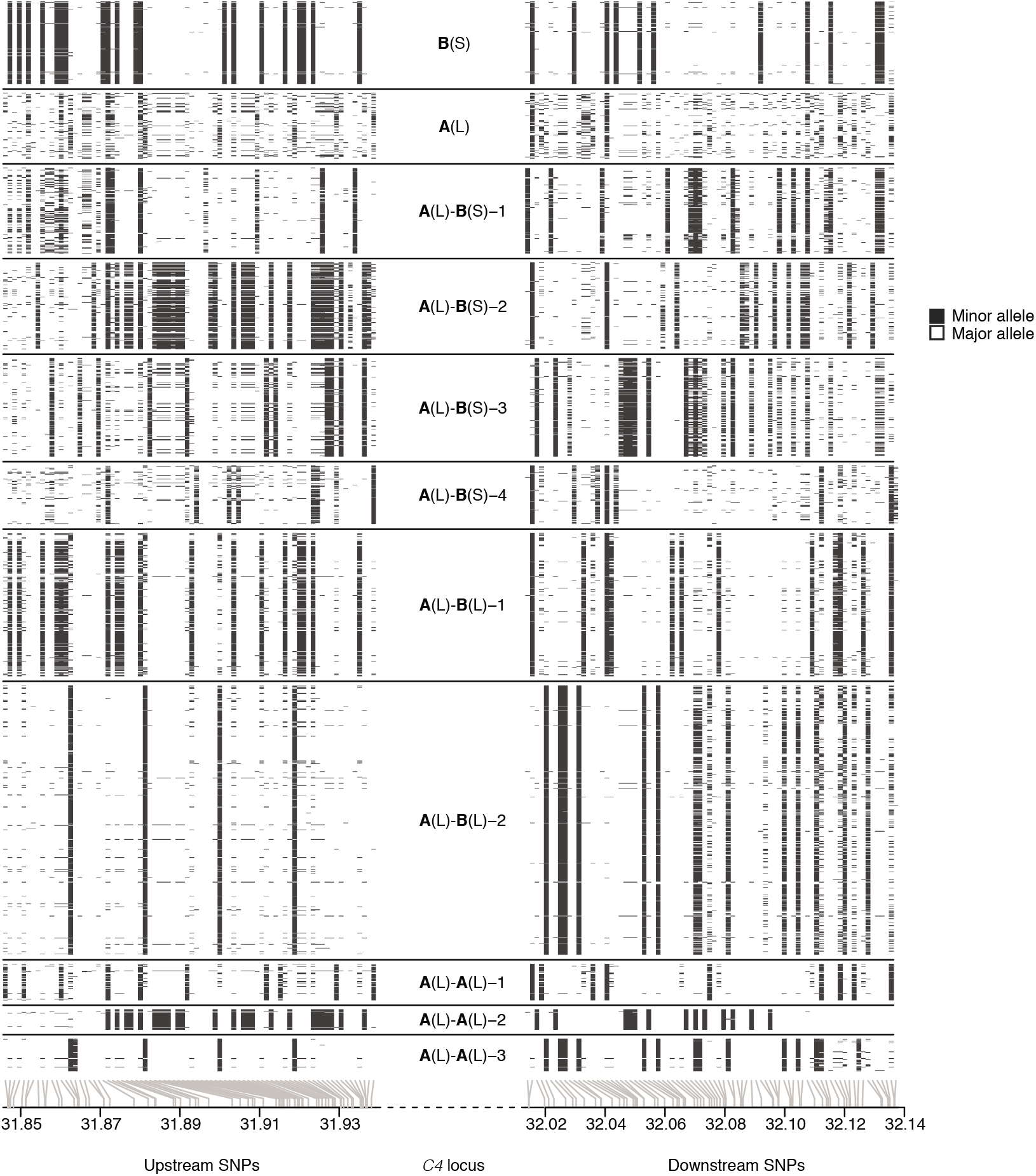
A panel of 2,530 reference haplotypes (created from whole-genome sequence data) containing *C4* alleles and SNPs across the MHC locus enables imputation of *C4* alleles into large-scale SNP data. The SNP haplotypes flanking each *C4* allele are shown as rows, with white and black representing the major and minor allele of each SNP as columns, respectively. Gray lines at the bottom indicate the physical location of each SNP along chromosome 6. The differences among the haplotypes are most pronounced closest to *C4* (toward the center of the plot), as historical recombination events in the flanking megabases will have caused the haplotypes to be less consistently distinct at greater genomic distances from *C4*. The patterns indicate that many combinations of *C4A* and *C4B* gene copy numbers have arisen recurrently on more than one SNP haplotype, a relationship that can be used in association analyses (**Fig. 2c**).

**Extended Data Figure 2.**
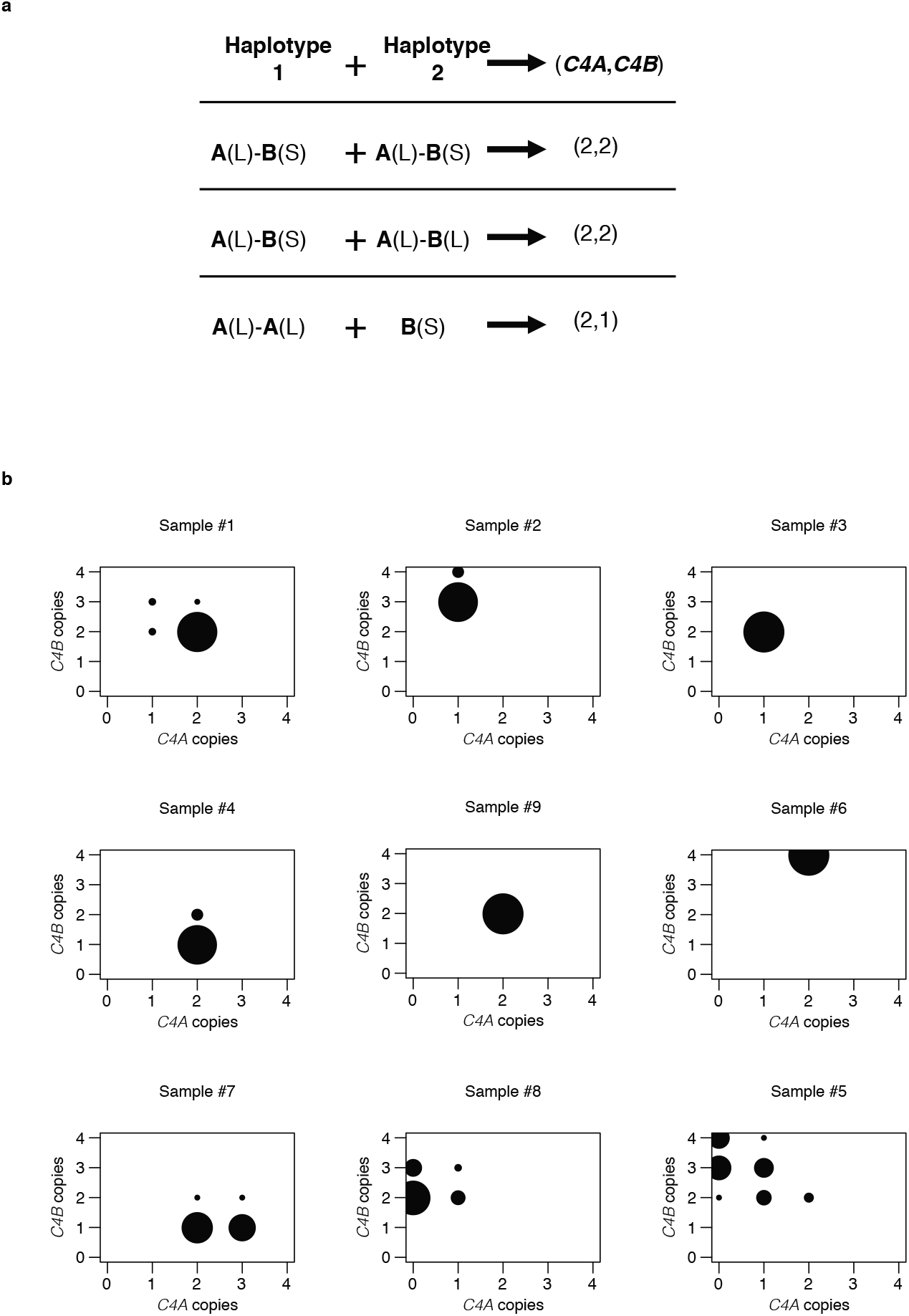
Aggregation of joint *C4A* and *C4B* genotypes probabilities per individual across imputed *C4* structural alleles for estimation of SLE risk for each combination. (a) An individual’s joint *C4A* and *C4B* gene copy number can be calculated by summing the *C4A* and *C4B* gene contents for each possible pair of two inherited alleles. Many pairings of possible inherited alleles result in the same joint *C4A* and *C4B* gene copy number. (b) Each individual’s *C4A* and *C4B* gene copy number was imputed from their SNP data, using the reference haplotypes summarized in **Extended Data Fig. 1**. For >95% of individuals (exemplified by samples 1–6 in the figure), this inference can be made with >90% certainty/confidence (the areas of the circles represent the posterior probability distribution over possible *C4A/C4B* gene copy numbers). For the remaining individuals (exemplified by samples 7–9 in the figure), greater statistical uncertainty persists about *C4* genotype. To account for this uncertainty, in downstream association analysis, all *C4* genotype assignments are handled as probabilistic gene dosages – analogous to the genotype dosages that are routinely used in large-scale genetic association studies that use imputation.

**Extended Data Figure 3.**
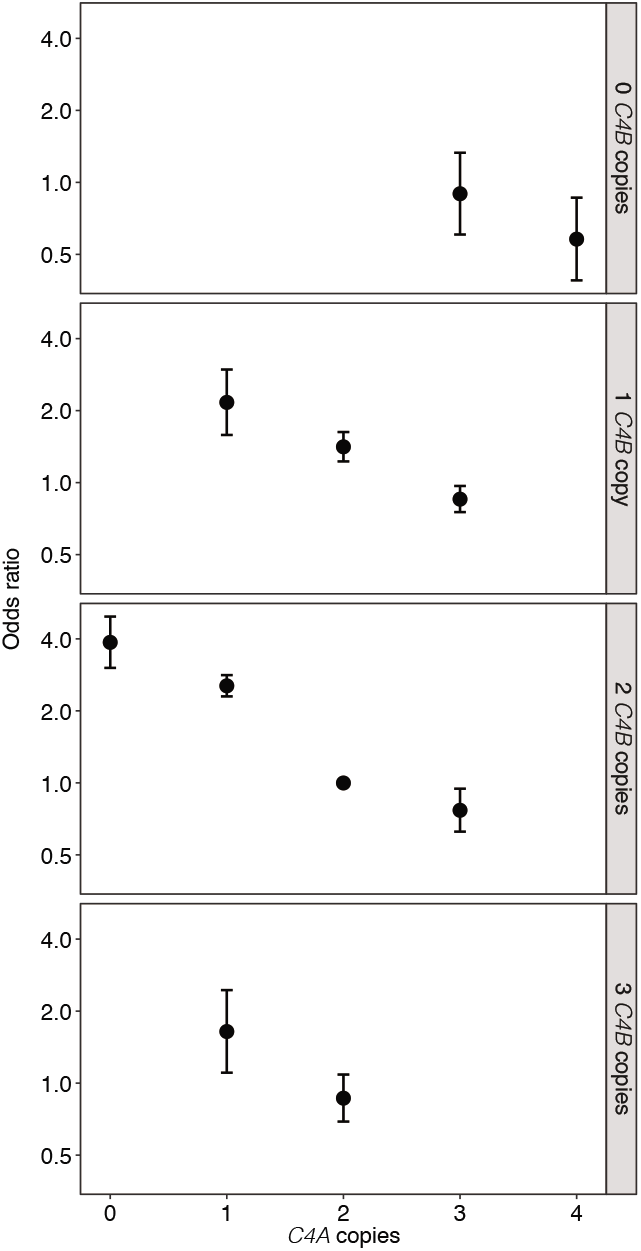
SLE odds ratios and confidence intervals for each combination of *C4A* and *C4B* gene copy number. Odds ratios and 95% confidence intervals underlying each of the *C4*-genotype risk estimates in **Fig. 2a** presented as a series of panels for each observed copy number of *C4B*, with increasing copy number of *C4A* for that *C4B* dosage (x-axis).

**Extended Data Figure 4.**
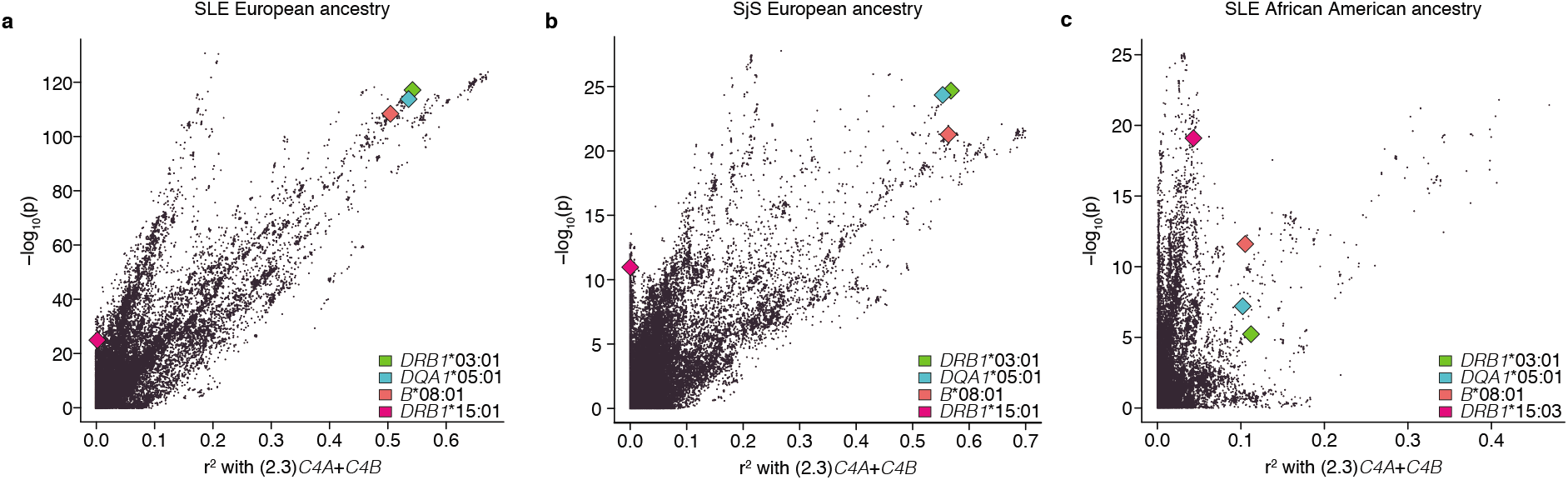
Relationship between association with SLE and linkage to *C4* for variants in the MHC region. (a) Relationship between SLE association [-log_10_(*p*), y-axis] and LD to the weighted *C4* risk score (x-axis) for genetic markers and imputed *HLA* alleles across the extended MHC locus. In this European-ancestry cohort, it is unclear (from this analysis alone) whether the association with the markers in the predominant ray of points (at a ~45° angle from the x-axis) is driven by variation at *C4* or by the long haplotype containing *DRB1*\*03:01 (green), *DQA1*\*05:01 (blue), and *B*\*08:01 (red). In addition, at least one independent association signal (a ray of points at a higher angle in the plot, with strong association signals and only weak LD-based correlation to *C4* and *DRB1*\*0301) with some LD to *DRB1*\*15:01 (maroon) is also present. (b) As in (a) but among the European-ancestry SjS cohort. Similar to SLE, it is unclear whether the effect is driven by variation at *C4* or linked *HLA* alleles, *DRB1*\*03:01 (green), *DQA1*\*05:01 (blue), and *B*\*08:01 (red). There is also an independent association signal with LD to *DRB1*\*15:01 (maroon). (c) Analysis of an African American SLE case–control cohort, in which LD in the MHC region is more limited, identified a set of markers that associate with SLE in proportion to their correlation with the *C4* composite risk score inferred from the earlier analysis of the European cohort, which itself associates with SLE at *p* < 10^−18^. No similar relationship is observed for *DRB1*\*03:01 and other alleles linked in European ancestry haplotypes. An independent association signal is also present in this cohort, more clearly in LD with the *DRB1*\*15:03 allele (maroon).

**Extended Data Figure 5.**
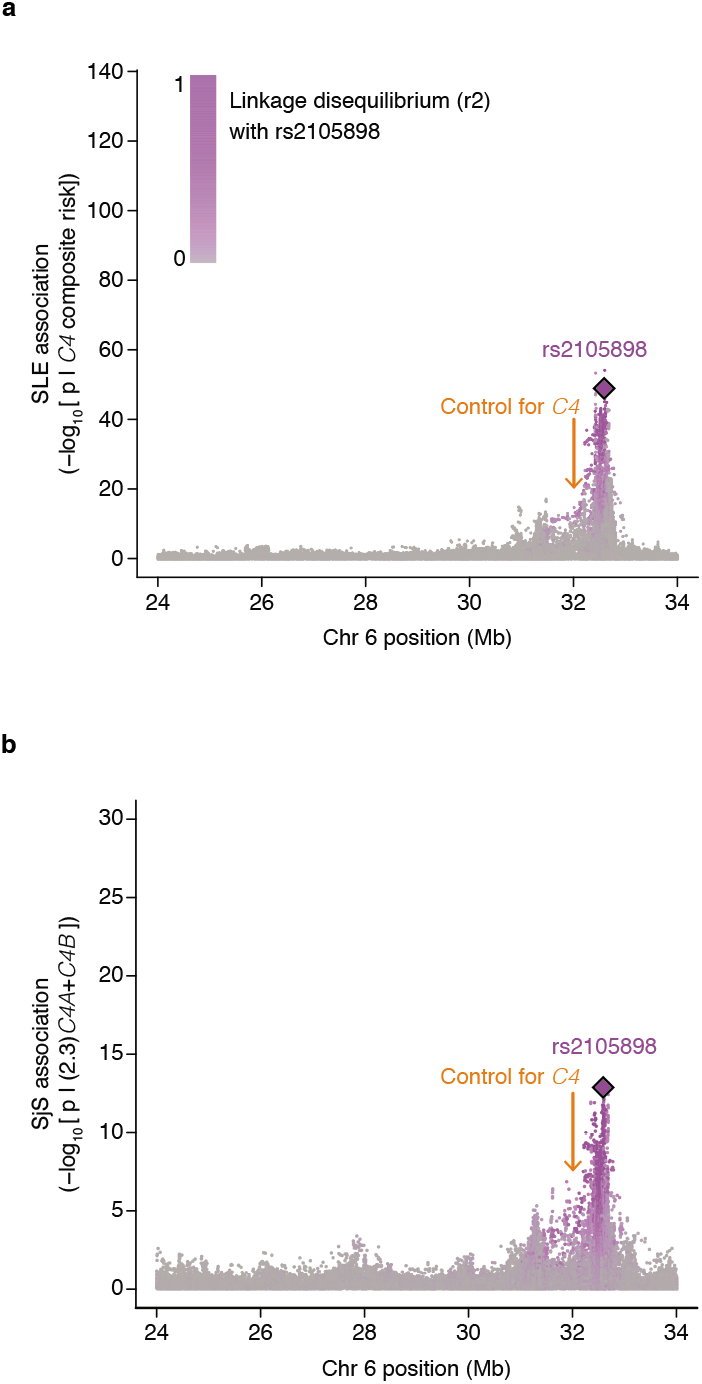
Conditional association analyses for genetic markers across the extended MHC locus within the European-ancestry cohort. (a) Association of SLE with genetic markers (SNPs and imputed *HLA* alleles) across the extended MHC locus within the European-ancestry cohort controlling for *C4* composite risk (weighted sum of risk associated with various combinations of *C4A* and *C4B*). Variants are shaded in purple by their LD with rs2105898, an independent association identified from trans-ancestral analyses. (b) As in (a), but in association with a European-ancestry SjS cohort. Here a simpler linear model of risk contributed by *C4A* and *C4B* was used instead of a weighted sum across all possible combinations.

**Extended Data Figure 6.**
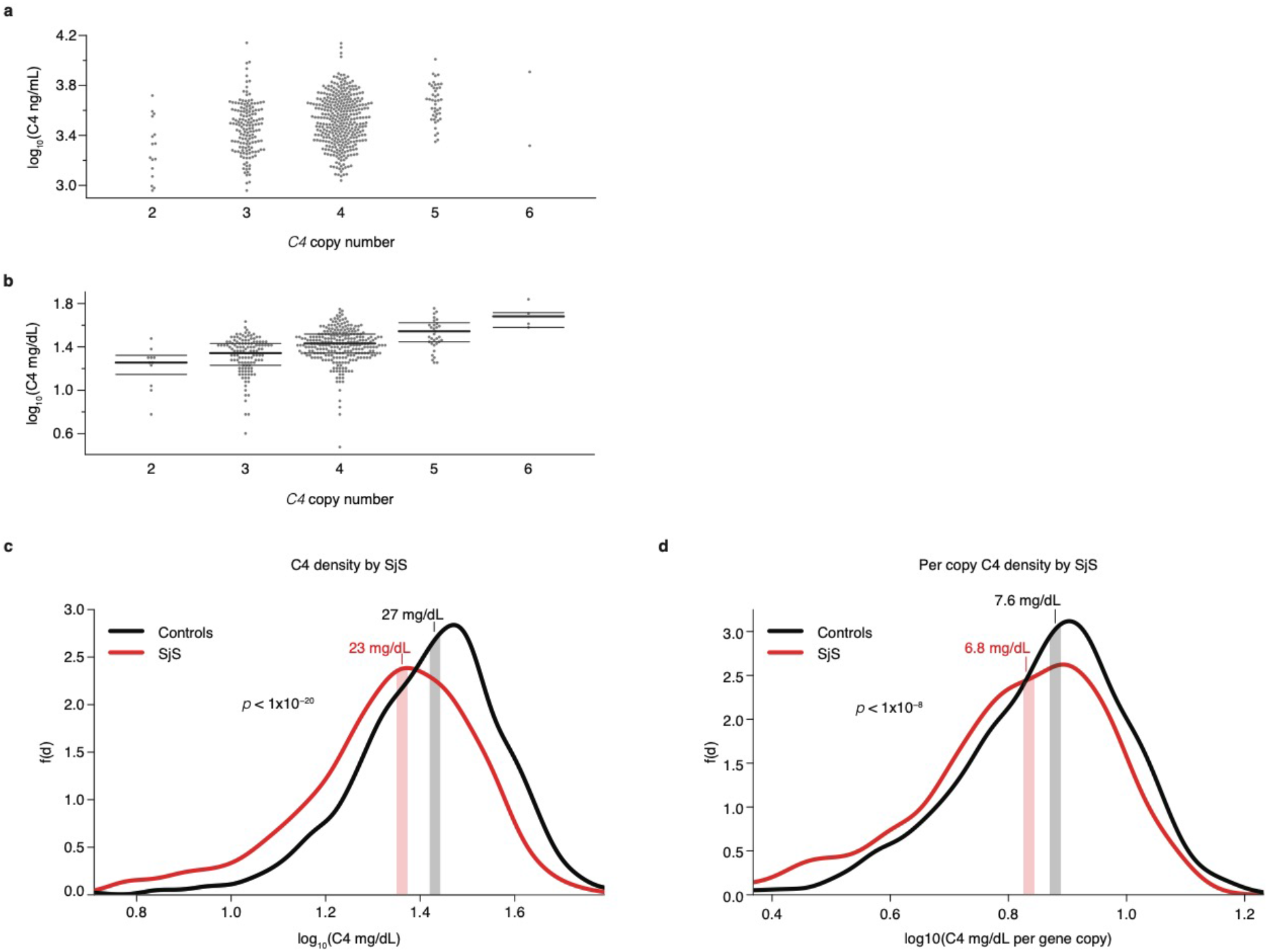
Correlation of C4 protein measurements (in cerebrospinal fluid and blood plasma) with imputed *C4* gene copy number. (a) Measurements of C4 protein in CSF obtained by ELISA are presented as log_10_(ng/mL) (y-axis) for each observed or imputed copy number of total *C4* (x-axis, here showing most likely copy number from imputation). Because *C4* gene copy number affects C4 protein levels so strongly, we normalized C4 protein measurements by *C4* gene copy number in subsequent analyses (**Fig. 4f**). (b) Measurements of C4 protein in blood plasma obtained by immunoturbidimetric assays are presented as log_10_(mg/dL) (y-axis) for each best-guess imputed copy number of total *C4* (x-axis). Because *C4* gene copy number affects C4 protein levels so strongly, we normalized C4 protein measurements by *C4* gene copy number in subsequent analyses (**Fig. 4h**). Due to the number of observations (*n* = 1,844 total), we downsampled to 500 points shown, but median and quartiles shown are for all individuals per C4 copy number. (c) C4 protein in blood plasma was measured in 670 individuals with SjS (red) and 1,151 individuals without SjS (black) and shown on a log_10_ scale (x-axis). Vertical stripes represent median levels for cases and controls separately. (d) As in (c), but concentrations are normalized to the number of *C4* gene copies in an individual’s genome and this per-copy amount is shown on a log_10_ scale (x-axis).

**Extended Data Figure 7.**
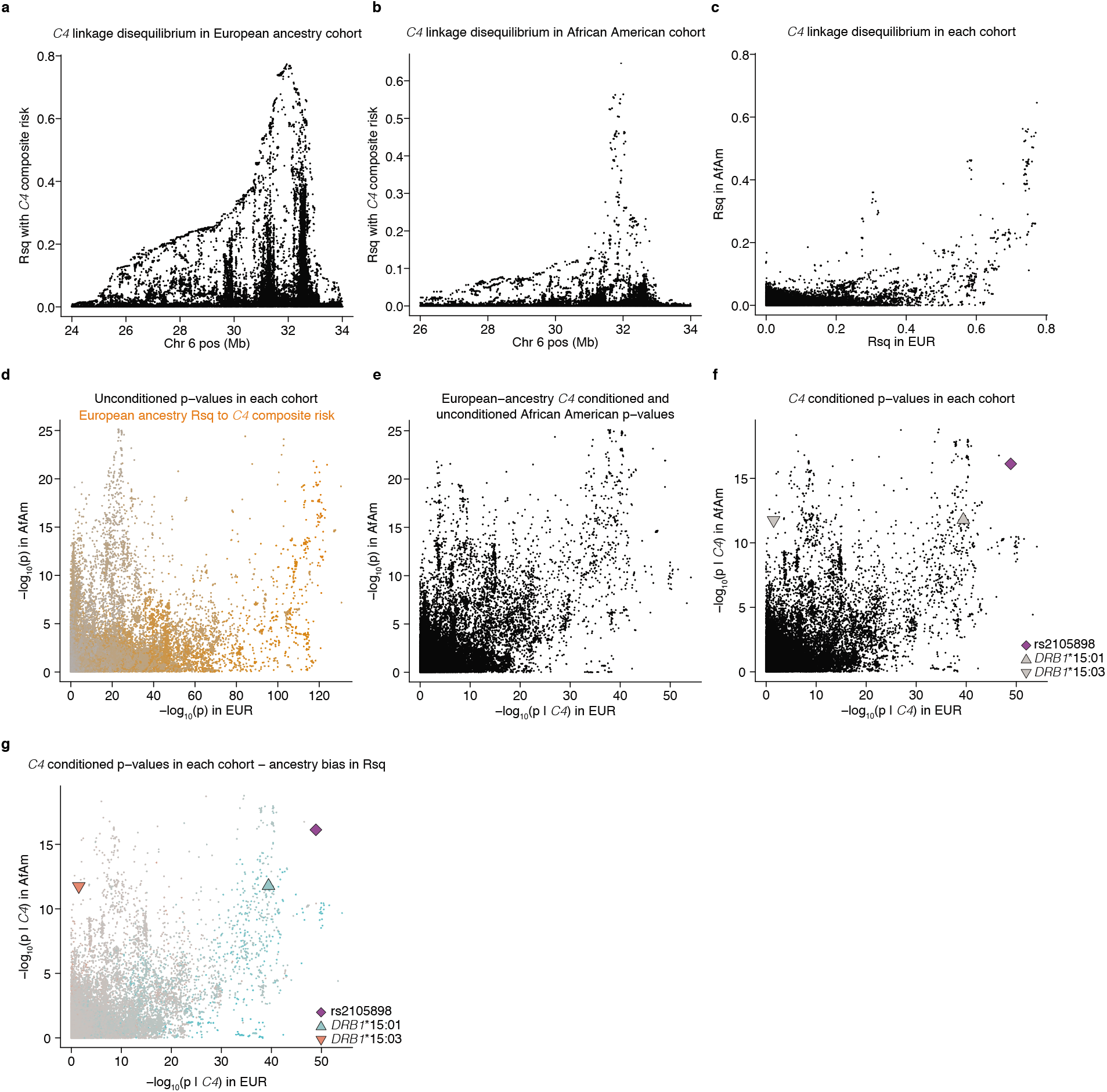
Concordance of trans-ancestral SLE risk association patterns across the MHC region largely a function of strong European LD between *C4* and nearby variants. (a) LD in European ancestry between the composite *C4* risk term (weighted sum of risk associated with various combinations of *C4A* and *C4B*) and variants in the MHC region as r^2^ (y-axis). (b) As in (a), but for African Americans. (c) LD for the same variants measured in European ancestry individuals (x-axis) and African Americans (y-axis). Note the abundance of variants that have greater LD with *C4* across European ancestry individuals, with several groups of variants that have similar LD in European ancestry individuals but exhibit a range of LD in African Americans. (d) Associations with SLE for the same variants in European ancestry cases and controls (x-axis) and African American cases and controls (y-axis). Variants are shaded in orange by their LD with *C4* in European ancestry individuals to highlight the effect of European-specific LD on shaping the discordant patterns of trans-ancestral associations with SLE risk in the MHC region. (e) As in (d), but controlling for the effect of *C4* in only European ancestry associations (x-axis). Note that this greatly aligns the patterns of association across the MHC region between European ancestry and African American cohorts. (f) As in (e), but controlling for the effect of *C4* in African American associations as well (y-axis). Note that this does not significantly affect the concordance seen in (e) due to the lack of broad LD relationships between *C4* and variants in the MHC region in African Americans. The independent signal, rs2105898, and *HLA* alleles, *DRB1*\*15:01 and *DRB1*\*15:03, are also highlighted. (g) As in (f), but with variants colored by whether they exhibit greater LD to rs2105898 in European ancestry individuals (blue) or African Americans (red). Note that the independent *DRB1*\*15:01 / *DRB1*\*15:03 association may be largely due to LD with rs2105898, with the relative strength of association for each in a particular cohort may be due to ancestry-specific LD with the haplotype defined by rs2105898. (DRB1*15:03 is largely an African-restricted allele, and *DRB1*\*15:01 may be picking up signal in African Americans during imputation – beyond the small fraction of admixed haplotypes – due to small dosages assigned by the classifier in haplotypes that likely have *DRB1*\*15:03.)

**Extended Data Figure 8.**
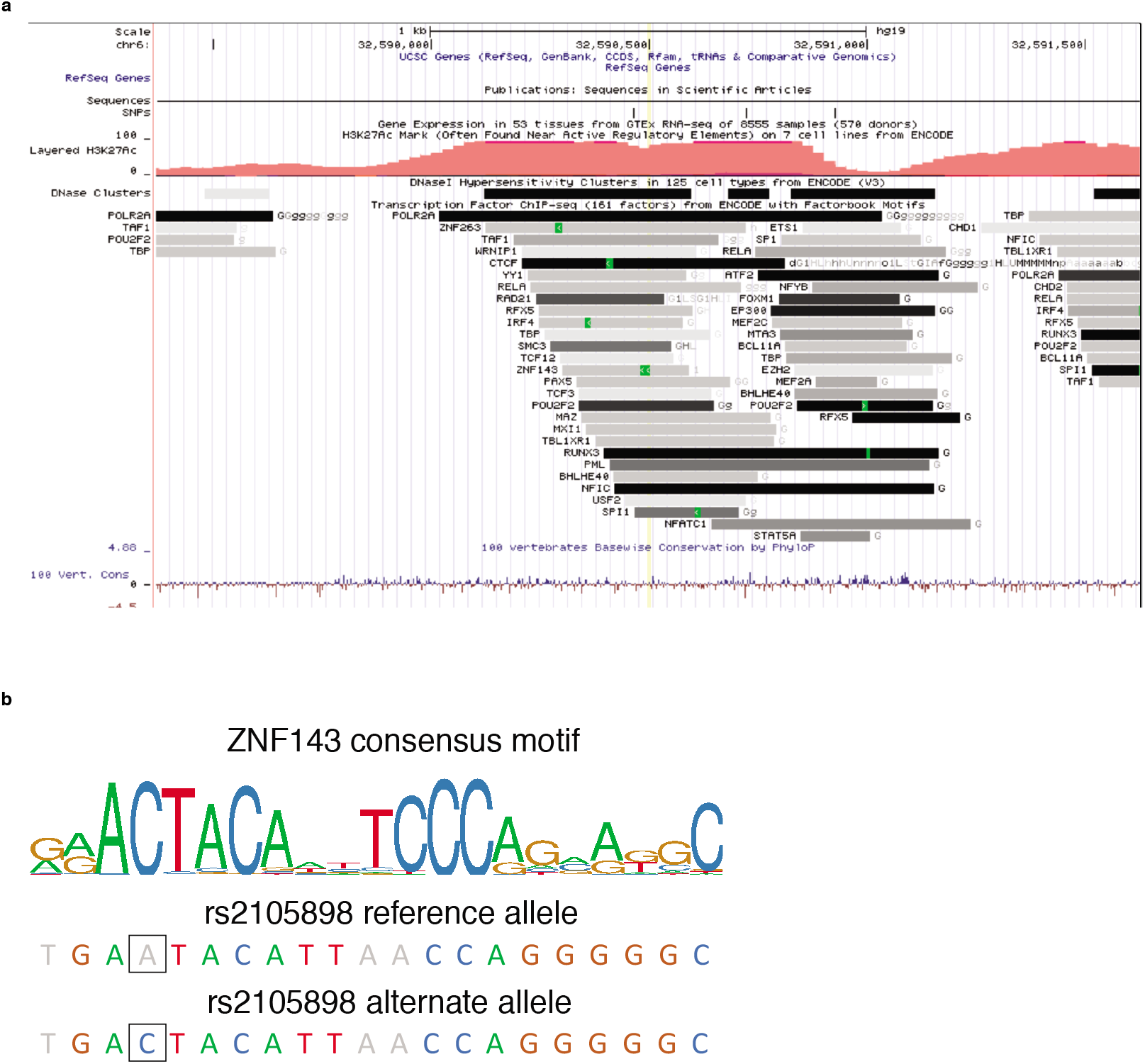
Effect of rs2105898 alleles on concordance with known ZNF143 binding motif in XL9 region. (a) Location of rs2105898 (yellow line at center) within the XL9 region, with relevant tracks showing overlapping histone marks and transcription factor binding peaks (from ENCODE^50^), visualized with the UCSC genome browser^63^. (b) ZNF143 consensus binding motif as a sequence logo, with the letters colored if the base is present in >5% of observed instances. The alleles of rs2105898 are indicated by outlined box surrounding the base.

**Extended Data Table 1.** Imputation accuracy for *C4* copy numbers in European ancestry and African American haplotypes. Accuracy was determined by cross-validation of the reference panel with directly-typed *C4* copy numbers from WGS data. Aggregated copy numbers imputed from each round of leaving 10 samples out were then correlated with the directly-typed measurements and reported as r^2^ for each type of copy number variation for European ancestry and African American members of the reference panel separately.

| **Gene copy number** | **Imputation accuracy ($r^2$)** |  |
| --- | --- | --- |
|  | **European ancestry** | **African Americans** |
| *C4* | 0.80 | 0.58 |
| *C4A* | 0.78 | 0.65 |
| *C4B* | 0.74 | 0.61 |
| *C4*-HERV | 0.91 | 0.76 |
| *2.3(C4A)+C4B* | 0.77 | 0.64 |

**Extended Data Table 2.**
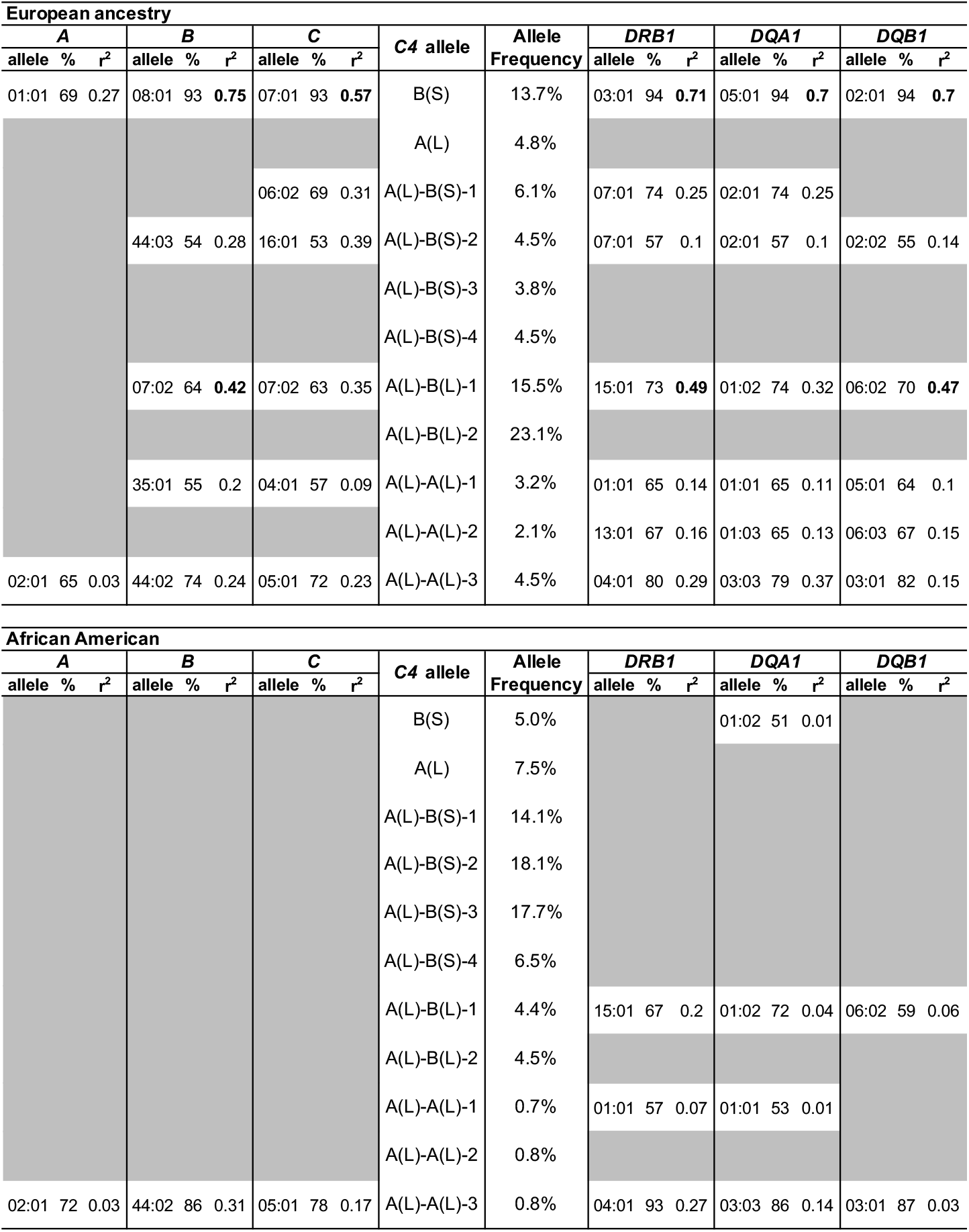
Frequency of common *C4* alleles and their linkage with *HLA* alleles in European ancestry and African American cohorts. For each common *C4* allele and *HLA* gene, the allele with highest LD (*r*^2^) is listed if present on more than half of the haplotypes with that *C4* allele (exact fraction in %). r^2^ values higher than 0.4 are highlighted to point out particularly strong *C4-HLA* allele pairings, such as for several with the *C4*-**B**(S) allele in European ancestry individuals. Some common *C4* alleles are further subdivided into distinct haplotypes used in imputation (and in **Fig. 2c**), as defined by shared alleles from variants flanking *C4*. Note that some alleles such as *C4*-**A**(L)-**A**(L)-3 are present at a frequency in African Americans that may solely reflect their presence on a fraction (~15-20%) of admixed haplotypes spanning this region, whereas others such as *C4*-**B**(S) are likely to also exist on African haplotypes – these differences between *C4* alleles are also reflected in the similarity of LD with *HLA* alleles to the corresponding row of the European ancestry section.

**Extended Data Table 3.** Logistic regression models of SLE risk against *C4* variation, *HLA* alleles, and/or rs2105898 in European ancestry and African American cohorts. Coefficients (beta, standard error) and p-values (as −log_10_(p)) for individual terms composing several relevant logistic regression models for predicting SLE risk that also include ancestry-specific covariates. For each model, the Akaike information criterion (AIC) and overall p-value (as determined by Chi-squared likelihood-ratio test) are given at the right end to indicate the relative strengths between similar models for each ancestry cohort.

| European ancestry |  |  |  |  |  |  |  |  |  |  |  |  |  |  |  |  |  |  |  |  |  |
| --- | --- | --- | --- | --- | --- | --- | --- | --- | --- | --- | --- | --- | --- | --- | --- | --- | --- | --- | --- | --- | --- |
| Model | C4 |  |  | C4A |  |  | C4B |  |  | DRB1*03:01 |  |  | B*08:01 |  |  | rs2105898 |  |  | AIC | LRT | -log<sub>10</sub>(p) |
|  | beta | se | -log<sub>10</sub>(p) | beta | se | -log<sub>10</sub>(p) | beta | se | -log<sub>10</sub>(p) | beta | se | -log<sub>10</sub>(p) | beta | se | -log<sub>10</sub>(p) | beta | se | -log<sub>10</sub>(p) |  |  |  |
| C4 | -0.55 | 0.027 | 92.7 |  |  |  |  |  |  |  |  |  |  |  |  |  |  |  | 22855.26 | 260.2 |  |
| C4A |  |  |  | -0.53 | 0.024 | 105.3 |  |  |  |  |  |  |  |  |  |  |  |  | 22790.05 | 274.3 |  |
| C4A+C4B |  |  |  | -0.62 | 0.028 | 112 | -0.27 | 0.037 | 12.3 |  |  |  |  |  |  |  |  |  | 22739.8 | 284.4 |  |
| DRB1*03:01 |  |  |  |  |  |  |  |  |  | 0.7 | 0.03 | 117.1 |  |  |  |  |  |  | 22748.33 | 283.3 |  |
| B*08:01 |  |  |  |  |  |  |  |  |  |  |  |  | 0.69 | 0.031 | 108.4 |  |  |  | 22790.65 | 274.2 |  |
| rs2105898 |  |  |  |  |  |  |  |  |  |  |  |  |  |  |  | -0.32 | 0.027 | 30.7 | 23153.86 | 195.5 |  |
| C4A + C4B + DRB1*03:01 |  |  |  | -0.35 | 0.041 | 17.2 | -0.11 | 0.041 | 2.3 | 0.4 | 0.046 | 17.5 |  |  |  |  |  |  | 22666.1 | 299.6 |  |
| C4A + C4B + B*08:01 |  |  |  | -0.41 | 0.039 | 24.6 | -0.17 | 0.039 | 4.7 |  |  |  | 0.35 | 0.044 | 14.4 |  |  |  | 22680.53 | 296.4 |  |
| C4A + C4B + rs2105898 |  |  |  | -0.67 | 0.028 | 122.8 | -0.32 | 0.038 | 16.4 |  |  |  |  |  |  | -0.38 | 0.028 | 41.1 | 22558.42 | 322.8 |  |

**Extended Data Table 3. Logistic regression models of SLE risk against C4 variation, HLA alleles, and/or rs2105898 in European ancestry and African American cohorts.**
| African American |  |  |  |  |  |  |  |  |  |  |  |  |  |  |  |  |  |  |  |  |  |
| --- | --- | --- | --- | --- | --- | --- | --- | --- | --- | --- | --- | --- | --- | --- | --- | --- | --- | --- | --- | --- | --- |
| Model | C4 |  |  | C4A |  |  | C4B |  |  | DRB1*03:01 |  |  | B*08:01 |  |  | rs2105898 |  |  | AIC | LRT | -log<sub>10</sub>(p) |
|  | beta | se | -log<sub>10</sub>(p) | beta | se | -log<sub>10</sub>(p) | beta | se | -log<sub>10</sub>(p) | beta | se | -log<sub>10</sub>(p) | beta | se | -log<sub>10</sub>(p) | beta | se | -log<sub>10</sub>(p) |  |  |  |
| C4 | -0.51 | 0.059 | 17.3 |  |  |  |  |  |  |  |  |  |  |  |  |  |  |  | 7358.65 | 19.7 |  |
| C4A |  |  |  | -0.43 | 0.062 | 11.2 |  |  |  |  |  |  |  |  |  |  |  |  | 7385.17 | 14 |  |
| C4A+C4B |  |  |  | -0.62 | 0.068 | 18.7 | -0.41 | 0.068 | 8.6 |  |  |  |  |  |  |  |  |  | 7351.45 | 20.9 |  |
| DRB1*03:01 |  |  |  |  |  |  |  |  |  | 0.41 | 0.091 | 5.2 |  |  |  |  |  |  | 7413.36 | 8 |  |
| B*08:01 |  |  |  |  |  |  |  |  |  |  |  |  | 0.78 | 0.11 | 11.6 |  |  |  | 7387.33 | 13.6 |  |
| rs2105898 |  |  |  |  |  |  |  |  |  |  |  |  |  |  |  | -0.46 | 0.047 | 21.9 | 7339.35 | 23.9 |  |
| C4A + C4B + DRB1*03:01 |  |  |  | -0.59 | 0.073 | 15 | -0.38 | 0.071 | 7.1 | 0.1 | 0.099 | 0.5 |  |  |  |  |  |  | 7352.34 | 20.4 |  |
| C4A + C4B + B*08:01 |  |  |  | -0.51 | 0.073 | 11.7 | -0.37 | 0.069 | 7.2 |  |  |  | 0.49 | 0.12 | 4.4 |  |  |  | 7337.24 | 23.6 |  |
| C4A + C4B + rs2105898 |  |  |  | -0.52 | 0.07 | 13.2 | -0.43 | 0.069 | 9.4 |  |  |  |  |  |  | -0.42 | 0.048 | 17.8 | 7277.78 | 36.2 |  |

## Methods

### Creation of a *C4* reference panel from whole-genome sequence data

We constructed a reference panel for imputation of *C4* structural haplotypes using whole-genome sequencing data for 1265 individuals from the Genomic Psychiatry Cohort^39^. The reference panel included individuals of diverse ancestry, including 765 Europeans, 250 African Americans, and 250 people of reported Latino ancestry.

We estimated the diploid *C4* copy number, and separately the diploid copy number of the contained HERV segment, using Genome STRiP^64^. Briefly, Genome STRiP carefully calibrates measurements of read depth across specific genomic segments of interest by estimating and normalizing away sample-specific technical effects such as the effect of GC content on read depth (estimated from the genome-wide data). To estimate *C4* copy number, we genotyped the segments 6:31948358–31981050 and 6:31981096–32013904 (hg19) for total copy number, but masked the intronic HERV segments that distinguish short (S) from long (L) *C4* gene isotypes. For the HERV region, we genotyped segments 6:31952461–31958829 and 6:31985199–31991567 (hg19) for total copy number. Across the 1,265 individuals, the resultant locus-specific copy-number estimates exhibited a strongly multi-modal distribution (**Fig. 1a**) from which individuals’ total *C4* copy numbers could be readily inferred.

We then estimated the ratio of *C4A* to *C4B* genes in each individual genome. To do this, we extracted reads mapping to the paralogous sequence variants that distinguish *C4A* from *C4B* (hg19 coordinates 6:31963859–31963876 and 6:31996597–31996614) in each individual, combining reads across the two sites. We included only reads that aligned to one of these segments in its entirety. We then counted the number of reads matching the canonical active site sequences for *C4A* (CCC TGT CCA GTG TTA GAC) and *C4B* (CTC TCT CCA GTG ATA CAT). We combined these counts with the likelihood estimates of diploid *C4* copy number (from Genome STRiP) to determine the maximum likelihood combination of *C4A* and *C4B* in each individual. We estimated the genotype quality of the *C4A* and *C4B* estimate from the likelihood ratio between the most likely and second most likely combinations.

To phase the *C4* haplotypes, we first used the GenerateHaploidCNVGenotypes utility in Genome STRiP to estimate haplotype-specific copy-number likelihoods for *C4* (total *C4* gene copy number), *C4A*, *C4B*, and HERV using the diploid likelihoods from the prior step as input. Default parameters for GenerateHaploidCNVGenotypes were used, plus -genotypeLikelihoodThreshold 0.0001. The output was then processed by the GenerateCNVHaplotypes utility in Genome STRiP to combine the multiple estimates into likelihood estimates for a set of unified structural alleles. GenerateCNVHaplotypes was run with default parameters, plus -defaultLogLikelihood −50, -unknownHaplotypeLikelihood −50, and - sampleHaplotypePriorLikelihood 2.0. The resultant VCF was phased using Beagle 4.1 (beagle_4.1_27Jul16.86a) in two steps: first, performing genotype refinement from the genotype likelihoods using the Beagle gtgl= and -maxlr=1000000 parameters, and then running Beagle again on the output file using gt= to complete the phasing.

Our previous work suggested that several *C4* structures segregate on different haplotypes, and probably arose by recurrent mutation on different haplotype backgrounds^14^. The GenerateCNVHaplotypes utility requires as input an enumerated set of structural alleles to assign to the samples in the reference cohort, including any structurally equivalent alleles, with distinct labels to mark them as independent, plus a list of samples to assign (with high likelihood) to specific labeled input alleles to disambiguate among these recurrent alleles. The selection of the set of structural alleles to be modeled, along with the labeling strategy, is important to our methodology and the performance of the reference panel. In the reference panel, each input allele represents a specific copy number structure and optionally includes a label that differentiates the allele from other independent alleles with equivalent structure. We use the notation <H_n_n_n_n_L> to identify each allele, where the four integers following the H are, respectively, the (redundant) haploid count of the total number of *C4* copies, *C4A* copies, *C4B* copies and HERV copies on the haplotype. For example, <H_2_1_1_1> was used to represent the “AL-BS” haplotype. The optional final label L is used to distinguish potentially recurrent haplotypes with otherwise equivalent structures (under the model) that should be treated as independent alleles for phasing and imputation.

To build the reference panel, we experimentally evaluated a large number of potential sets of structural alleles and methods for assigning labels to potentially recurrent alleles. For each evaluation, we built a reference panel using the 1265 reference samples, and then evaluated the performance of the panel via cross-validation, leaving out 10 different samples in each trial (5 samples in the last trial) and imputing the missing samples from the remaining samples in the panel. The imputed results for all 1265 samples were then compared to the original diploid copy number estimates to evaluate the performance of each candidate reference panel (**Extended Data Table 1**).

Using this procedure, we selected a final panel for downstream analysis that used a set of 29 structural alleles representing 16 distinct allelic structures (as listed in the reference panel VCF file). Each allele contained from one to three copies of *C4*. Three allelic structures (AL-BS, AL-BL, and AL-AL) were represented as a set of independently labeled alleles with 9, 3, and 4 labels, respectively.

To identify the number of labels to use on the different alleles and the samples to “seed” the alleles, we generated “spider plots” of the *C4* locus based on initial phasing experiments run without labeled alleles, and then clustered the resulting haplotypes in two dimensions based on the Y-coordinate distance between the haplotypes on the left and right sides of the spider plot. Clustering was based on visualizing the clusters (**Extended Data Fig. 1**) and then manually choosing both the number of clusters (labels) to assign and a set of confidently assigned haplotypes to use to “seed” the clusters in GenerateCNVHaplotypes. This procedure was iterated multiple times using cross-validation, as described above, to evaluate the imputation performance of each candidate labeling strategy.

Within the data set used to build the reference panel, there is evidence for individuals carrying seven or more diploid copies of *C4*, which implies the existence of (rare) alleles with four or more copies of *C4*. In our experiments, attempting to add additional haplotypes to model these rare four-copy alleles reduced overall imputation performance. Consequently, we conducted all downstream analyses using a reference panel that models only alleles with up to three copies of *C4*. In the future, larger reference panels might benefit from modeling these rare four-copy alleles.

The reference panel will be available in dbGaP (accession # pending) with broad permission for research use.

### Genetic data for SLE

For analysis of systemic lupus erythematosus (SLE), collection and genotyping of the European-ancestry cohort (6,748 cases, 11,516 controls, genotyped by ImmunoChip) as previously described^7^. Collection and genotyping of the African-American cohort (1,494 cases, 5,908 controls, genotyped by OmniExpress) as previously described^11^.

### Genetic data for SjS

For analysis of Sjogren’s syndrome (SjS), collection and genotyping of the European-ancestry cohort (673 cases, 1,153 controls, genotyped by Omni2.5) as previously described^47^ and available in dbGaP under study accession number phs000672.v1.p1.

### Genetic data for schizophrenia

The schizophrenia analysis made use of genotype data from 40 cohorts of European ancestry (28,799 cases, 35,986 controls) made available by the Psychiatric Genetics Consortium (PGC) as previously described^62^. Genotyping chips used for each cohort are listed in Supplementary Table 3 of that study.

### Imputation of *C4* alleles

The reference haplotypes described above were used to extend the SLE, SjS, or schizophrenia cohort SNP genotypes by imputation. SNP data in VCF format were used as input for Beagle v4.1^65,66^ for imputation of *C4* as a multi-allelic variant. Within the Beagle pipeline, the reference panel was first converted to bref format. From the cohort SNP genotypes, we used only those SNPs from the MHC region (chr6:24-34 Mb on hg19) that were also in the haplotype reference panel. We used the conform-gt tool to perform strand-flipping and filtering of specific SNPs for which strand remained ambiguous. Beagle was run using default parameters with two key exceptions: we used the GRCh37 PLINK recombination map, and we set the output to include genotype probability (i.e., GP field in VCF) for correct downstream probabilistic estimation of *C4A* and *C4B* joint dosages.

### Imputation of *HLA* alleles

For *HLA* allele imputation, sample genotypes were used as input for the R package HIBAG^67^. For both European ancestry and African American cohorts, publicly available multi-ethnic reference panels generated for the most appropriate genotyping chip (i.e. Immunochip for European ancestry SLE cohort, Omni 2.5 for European ancestry SjS cohort, and OmniExpress for African American SLE cohort) were used^68^. Default parameters were used for all settings. All class I and class II *HLA* genes were imputed. Output haplotype posterior probabilities were summed per allele to yield diploid dosages for each individual.

### Associating single and joint *C4* structural allele dosages to SLE and SjS in European ancestry individuals

The analysis described above yields dosage estimates for each of the common C4 structural haplotypes (e.g., AL-BS, AL-AL, etc.) for each genome in each cohort. In addition to performing association analysis on these structures (**Fig 2c**), we also performed association analysis on the dosages of each underlying *C4* gene isotype (i.e. *C4A*, *C4B*, *C4L*, and *C4S*). These dosages were computed from the allelic dosage (DS) field of the imputation output VCF simply by multiplying the dosage of a *C4* structural haplotype by the number of copies of each *C4* isotype that haplotype contains (e.g., AL-BL contains one *C4A* gene and one *C4B* gene).

*C4* isotype dosages were then tested for disease association by logistic regression, with the inclusion of four available ancestry covariates derived from genome-wide principal component analysis (PCA) as additional independent variables, PC_c_,

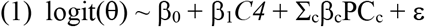

where θ=E[SLE|**X**]. For SjS, the model instead included two available multiethnic ancestry covariates from dbGaP that correlated strongly with European-specific ancestry covariates (specifically, PC5 and PC7) and smoking status as independent variables. Coefficients for relative weighting of *C4A* and *C4B* dosages were obtained from a joint logistic regression,

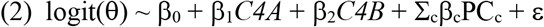

The values per individual of β_1_*C4A* + β_2_*C4B* were used as a combined *C4* risk term for estimating both association strength (**Fig. 2b**) as well as evaluating the relationship between the strength of nearby variants’ association with SLE or SjS and linkage with *C4* variation (**Extended Data Fig. 5a,b**).

Joint dosages of *C4A* and *C4B* for each individual in the same cohort were estimated by summing across their genotype probabilities of paired structural alleles that encode for the same diploid copy numbers of both *C4A* and *C4B* (**Extended Data Fig. 2a,b**). For each individual/genome, this yields a joint dosage distribution of *C4A* and *C4B* gene copy number, reflecting any possible imputed haplotype-level dosages with nonzero probability. Joint dosages for *C4A* and *C4B* diploid copy numbers were tested for association with SLE in a joint model with the same ancestry covariates (**Fig. 2a**),

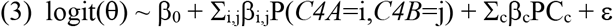

### Calculation of composite *C4* risk for SLE

Because SLE risk strongly associated with *C4A* and *C4B* copy numbers (**Fig. 2a**) in a manner that can be approximated as – but is not necessarily linear or independent – a composite *C4* risk score was derived by taking the weighted sum of joint *C4A* and *C4B* dosages multiplied by the corresponding effect sizes from the aforementioned model of the joint *C4A* and *C4B* diploid copy numbers. The weights for calculating this composite *C4* risk term were computed from the data from the European ancestry cohort, and then applied unchanged to analysis of the African American cohort.

### Associations of variants across the MHC region to SLE and SjS

Genotypes for non-array SNPs were imputed with IMPUTE2 using the 1000 Genomes reference panel; separate analyses were performed for the European-ancestry and African American cohorts. Unless otherwise stated, all subsequent SLE analyses were performed identically for both European ancestry and African American cohorts. Dosage of each variant, v_i_, was tested for association with SLE or SjS in a logistic regression including available ancestry covariates (and smoking status for SjS) first alone (**Fig. 2b, d**),

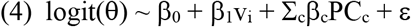

then with *C4* composite risk (**Extended Data Fig. 6a**),

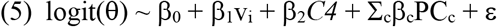

and finally with *C4* composite risk and rs2105898 dosage (**Extended Data Fig. 6b**),

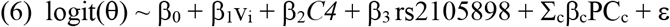

where θ=E[SLE|**X**]. For SjS, the simpler weighted (2.3)*C4A*+*C4B* model was used instead of composite risk term, as the cohort’s size gave poor precision to estimates of risk for many joint (*C4A*, *C4B*) copy numbers (**Extended Data Fig. 6c, d**). The Pearson correlation between the *C4* composite risk term and each other variant was computed and squared (r^2^) to yield a measure of linkage disequilibrium between *C4* composite risk and that variant in that cohort.

### Association analyses for specific *C4* structural alleles

The *C4* structural haplotypes were tested for association with disease (**Fig. 2c, 3a**) in a joint logistic regression that included (i) terms for dosages of the five most common *C4* structural haplotypes (AL-BS, AL-BL, AL-AL, BS, and AL), (ii) (for SLE and SjS) rs2105898 genotype, and (iii) ancestry covariates and (for SjS) smoking status,

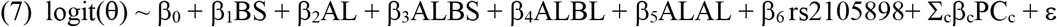

where θ=E[SLE|**X**]. Several of these common *C4* structural alleles arose multiple times on distinct haplotypes; we term the set of haplotypes in which such a common allele appeared as “haplogroups”. The haplogroups can be further tested in a logistic regression model in which the structural allele appearing in all member haplotypes is instead encoded as dosages for each of the SNP haplotypes in which it appears. These association analyses (**Fig. 2c**) were performed as in (6), with structural allele dosages for ALBS, ALBL, and ALAL replaced by multiple terms for each distinct haplotype.

To delineate the relationship between *C4*-BS and *DRB1*\*03:01 alleles – which are highly linked in European ancestry haplotypes – allelic dosages per individual in the African American SLE cohort were rounded to yield the most likely integer dosage for each. Although genotype dosages for each are reported by BEAGLE and HIBAG respectively, probabilities per haplotype are not linked and multiplying possible diploid dosages could yield incorrect non-zero joint dosages. Joint genotypes were tested as individual terms in a logistic regression model,

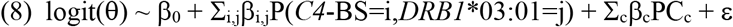

### Sex-stratified associations of *C4* structural alleles and other variants with SLE, SjS, and schizophrenia (Fig. 4a-d)

Determination of an effect from sex on the contribution of overall *C4* variation to risk for each disorder was done by including an interaction term between sex and *C4*; ie. (2.3)C4A+C4B for SLE and SjS and estimated C4A expression for schizophrenia:

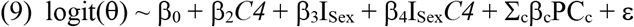

Each variant in the MHC region was tested for association with among European ancestry cases and cohorts in a logistic regression as in models (4)–(6) using only male cases and controls, and then separately using only female cases and controls (**Fig. 4c-e**). Likewise, allelic series analyses were performed as in (7), but in separate models for men and women (**Fig. 4a, b**).

To assess the relationship between sex bias in the risk associated with a variant and linkage to *C4* composite risk (as non-negative r^2^), male and female log-odds were multiplied by the sign of the Pearson correlation between that variant and *C4* composite risk before taking the difference.

### Analyses of cerebrospinal fluid

Cerebrospinal fluid (CSF) from healthy individuals was obtained from two research panels. The first panel, consisting of 533 donors (327 male, 126 female) from hospitals around Utrecht, Netherlands, was described previously^69,70^. The donors were generally healthy research participants undergoing spinal anesthesia for minor elective surgery. The same donors were previously genotyped using the Illumina Omni SNP array. To estimate *C4* copy numbers, we used SNPs from the MHC region (chr6:24-34 Mb on hg19) as input for *C4* allele imputation with Beagle, as described above in **Imputation of *C4* alleles**.

The second CSF panel sampled specimens from 56 donors (14 male, 42 female) from Brigham and Women’s Hospital (BWH; Boston, MA, USA) under a protocol approved by the institutional review board at BWH (IRB protocol ID no. 1999P010911). These samples were originally obtained to exclude the possibility of infection, and clinical analyses had revealed no evidence of infection. Donors ranged in age from 18 to 64 years old. Blood samples from the same individuals were used for extraction of genomic DNA, and *C4* gene copy number was measured by droplet digital PCR (ddPCR) as previously described^14^. Samples were excluded from measurements if they lacked *C4* genotypes, sex information, or contained visible blood contamination.

C4 measurements were performed by sandwich ELISA of 1:400 dilutions of the original CSF sample using goat anti-sera against human C4 as the capture antibody (Quidel, A305), FITC-conjugated polyclonal rabbit anti-human C4c as the detection antibody (Dako, F016902-2), and alkaline phosphatase–conjugated polyclonal goat anti-rabbit IgG as the secondary antibody (Abcam, ab97048). C3 measurements were performed using the human complement C3 ELISA kit (Abcam, ab108823).

Because *C4* gene copy number had a large and proportional effect on C4 protein concentration in these CSF samples (**Extended Data Fig. 7a**), we corrected for *C4* gene copy number in our analysis of relationship between sex and C4 protein concentration, by normalizing the ratio of C4 protein (in CSF) to *C4* gene copies (in genome). Therefore, these analyses included only samples for which DNA was available or *C4* was successfully imputed. In total, 495 (332 male, 163 female) C4 and 304 (179 male, 125 female) C3 concentrations were obtained across both cohorts. Log-concentrations of C3 (ng/mL) and C4 (ng/[mL, per *C4* gene copy number]) protein were then used separately in linear regression models to estimate a sex-unbiased cohort-specific offset for each protein,

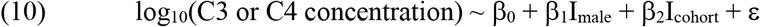

to be applied to all concentrations for that protein. Estimation of average measurements by age for each sex was done by local polynomial regression smoothing (LOESS). To evaluate the significance of sex effects, we used these cohort-corrected concentrations estimates and analyzed them with the non-parametric unsigned Mann-Whitney rank–sum test comparing concentration distributions for males and females.

### Analyses of blood plasma

Blood plasma was collected and immunoturbidimetric measurements of C3 and C4 protein in 1,844 individuals (182 men, 1662 women) by Sjögren’s International Collaborative Clinical Alliance (SICCA) from individuals with and without SjS as previously described^71^. *C4* copy numbers for these individuals were previously imputed for use in logistic regression of SjS risk. As *C4* copy number has an effect on measured C4 protein similar to CSF (**Extended Data Fig. 7b**), we normalized C4 levels to them in all following analyses. Estimation of average measurements by age for each sex was done by local polynomial regression smoothing (LOESS) on log-concentrations of C3 (mg/dL) and C4 (mg/[dL, per *C4* gene copy number]) protein. To evaluate the significance of sex bias within age ranges displaying the greatest difference (informed by LOESS), we analyzed individuals in these bins with the non-parametric unsigned Mann-Whitney rank-sum test comparing concentration distributions for males and females.

Difference in C4 protein levels between individual with and without SjS was done by performing a non-parametric unsigned Mann-Whitney rank–sum test on C4 protein levels normalized to *C4* genomic copy number (**Extended Data Fig. 7c**).

## Acknowledgements

This work was supported by the National Human Genome Research Institute (HG006855), the National Institute of Mental Health (MH112491, MH105641, MH105653), and the Stanley Center for Psychiatric Research. In addition, this work was supported by the Biomedical Research Centre based at Guy’s and St Thomas’ NHS Foundation Trust in partnership with King’s College London (D.L.M., P.T., and T.J.V.). We thank Christina Usher for contributions to the figures and manuscript text, Marta Florio for suggestions regarding figure display, and David Curtis and Chris Patil for suggestions on the manuscript.

## Author Contributions

S.A.M., N.K., and A.S. conceived the genetic studies. M.T.P., C.N.P., and M.B. collected and contributed whole-genome sequence data for the Genomic Psychiatry Cohort. R.E.H. and C.W.W. genotyped *C4* structural variation in the Genomic Psychiatry Cohort and optimized variant selection for use as a reference panel in the imputation of *C4* variation into lupus and schizophrenia cohorts (Fig. 1 and Extended Data Fig. 1). T.J.V., R.R.G., L.A.C., C.D.L., R.P.K., J.B.H., K.M.K., D.L.M., and P.T. contributed genotype data and imputation of non-C4 variation for analysis of SLE cohorts. K.E.T. and L.A.C. contributed genotype and phenotype data along with imputation of non-C4 variation for analysis of the SjS cohort. Investigators in the Schizophrenia Working Group of the Psychiatric Genomics Consortium collected and phenotyped cohorts and contributed genotype data for analysis of schizophrenia cohorts. N.K did the imputation and association analyses (Fig. 2, 3, 4a-e, and 4h, i and Extended Data Fig. 2–5, 6b-d, 7, and 8). T.J.V., R.R.G., and D.L.M. provided valuable advice on the analysis and interpretation of SLE association results. R.A.O. and L.M.O.L collected and provided CSF samples composing the group from Utrecht, Netherlands. C.E.S. collected and provided CSF samples composing the Brigham & Women’s Hospital group. H.d.R and K.T. performed the C4 and C3 immunoassay experiments on CSF samples (Fig. 4f, g and Extended Data Fig. 6a). S.A.M and N.K. wrote the manuscript with contributions from all authors.

