## Supplemental Note for "Complement component 4 genes contribute sex-specific vulnerability in diverse illnesses"

**Contents:**

Supplementary Note - Fine mapping of an independent association signal in the MHC class II region 1

Full list of collaborators from the PGC Schizophrenia Working Group 4

### Supplementary Note - Fine mapping of an independent association signal in the MHC class II region

#### Linkage disequilibrium of *C4* variation to other MHC variants differs by ancestry

The linkage-disequilibrium (LD) relationships of *C4* variation to other genetic variation in the MHC locus differ greatly in magnitude and pattern between European-ancestry and African American cohorts. For example, **Extended Data Fig. 7a, b** shows the LD-correlation ($r^2$) of SNPs across the MHC locus to the composite estimate of C4-derived SLE risk employed in this study. (Other *C4* features, such as total *C4* gene copy number, also exhibit strikingly different correlations with genetic markers between the two populations.). Most notably, LD in European ancestry is widespread across the extended MHC locus (**Extended Data Fig. 7a**) – and particularly strong in the nearby MHC class II region (32-33 Mb) – while strong LD in African Americans is localized primarily to a much-smaller region immediately flanking the *C4* genes (**Extended Data Fig. 7b**).

A direct comparison of the two population-specific LD patterns confirms that nearly all variants with LD to *C4* variation have greater LD in European-ancestry than in African American population sample, where only a small subset of European ancestry-linked alleles have similar or lower levels of linkage in African Americans (**Extended Data Fig. 7c**).

#### Initial (*C4*-naïve) association analysis produces divergent MHC association results in European-ancestry and African American cohorts

Unconditional (*C4*-naïve) association analysis of SLE of each variant in the MHC locus exhibits little correlation between European-ancestry and African American cohorts (**Extended Data Fig. 7d**). Of course, this could result from multiple population-specific variants or even population-specific biology, but in general, more-parsimonious explanations are both strongly preferred, and more strongly constrained and testable by available data.

Of course, *C4* alleles have both strong allele-frequency and LD differences between these populations (**Extended Data Table 1**) and therefore could be a potential contributor to these differences in **Extended Data Fig. 7d**.

To appreciate this possibility, the points in **Extended Data Fig. 7d** are colored orange in proportion to their European-ancestry LD ($r^2$) to *C4* composite risk. This highlights the strong effect that *C4* alleles would be likely to have in shaping the relative association strengths of genetic markers throughout the MHC locus.

#### Conditioning on *C4* composite risk in European-ancestry cohort only

Considering *C4* in the above analysis begins to strongly align the association signals in Europeans and African Americans.

If, beginning with the European-ancestry cohort, we now consider SNPs not in a naïve association analysis but in a joint association analysis together with *C4* (i.e. with *C4* genetic risk as a covariate), then the association statistics for variants in the two cohorts begin to align with each other more strongly (**Extended Data Fig. 7e**).

Adjusting the association statistics for the African-American cohort analysis to account for *C4* effects changes the overall pattern more modestly (**Extended Data Fig. 7f**). (This likely reflects reduced LD to

*C4* alleles among African Americans, and reduced *C4* variation among African Americans relative to Europeans.)

(Population-specific *HLA* alleles (*DRB1**15:01 and *DRB1**15:03) have been proposed as potential explanations for the apparently divergent association signals across European ancestry and African American populations. In **Extended Data Fig. 7f**, these variants are shown with grey triangles.)

Much of the population differences in SLE association pattern (that remain after controlling for *C4*) appear to be explained by differences in LD patterns between populations. In the same plots, coloring the variants by European or African American LD ($r^2$) to rs2105898 reveals that the variants with higher relative associations in the European ancestry cohort (lower right in below plots) generally have higher LD to rs2105898 in that cohort. (This includes the European ancestry-specific SLE association to the *HLA-DRB1**15:01 allele). Few variants have higher LD to rs2105898 in African Americans – though one such variant is the *HLA-DRB1**15:03 allele, which has previously been reported to associate to SLE specifically in African Americans.

Much of the remaining differences in association pattern appear to be explained by differences in LD patterns between populations; in **Extended Data Fig. 7g**, blueness represents greater LD to rs2105898 in the European-ancestry cohort (relative to LD among African Americans) and redness represents greater LD in the African American cohort (relative to LD among Europeans). Notably, the many variants with relatively stronger association signals among Europeans (including *DRB1**1501) exhibit stronger LD to rs2105898 among Europeans, while select variants with relatively stronger association signals among African Americans (including *DRB1**1503) exhibit stronger LD to rs2105898 among African Americans.

This analysis also indicates that while much, if not all, of the European ancestry-specific association after controlling for *C4* composite risk can be accounted for by European ancestry-specific LD to rs2105898, this is not true for African Americans, who may harbor at least one additional, independent genetic effect not explained by the above analysis.

**The *C4*-independent association signal comprising rs2105898 and another linked variant defines strong pan-tissue expression QTLs for *HLA* Class II genes**

Although rs2105898 was the top variant associated between cohorts in analysis controlling for *C4*, there is one other variant (rs9271513) in high ($r^2$>0.9) LD across both populations that should be considered together as a haplotype.

As mentioned in the main text, we found that rs2105898 (and the highly LD-correlated variant) are significant eQTLs for 171 gene-tissue associations, largely comprised of significant associations for 7 *HLA* Class II genes (*HLA-DRB1, HLA-DRB5, HLA-DRB6, HLA-DQA1, HLA-DQA2, HLA-DQB1, HLA-DQB2*) in almost every tissue sampled by the GTEx Consortium[33].

**The rs2105898 haplotype affects XL9 hotspot of active chromatin and transcription factor binding**

rs2105898 and the variant with which it is strong LD in both European and African American populations define a haplotype which is the effective unit of genetic association. rs2105898 in particular lies within multiple histone marks that are associated with active enhancers (6 tissues), in the XL9 region of open chromatin (15 tissues), and under ChIP-seq binding peaks for 19 transcription factors (**Extended Data Fig. 8a**, data from the ENCODE project[31]).

**rs2105898 disrupts a binding site for the ZNF143 transcription factor**

We identified transcription factors whose binding motif is significantly affected by rs2105898 allele. The strongest hit (ZNF143) is also among the transcription factors that have been determined by ChIP-seq analysis (from the ENCODE project) to bind to DNA sequence at rs2105898 (**Extended Data Fig. 8b)**. ZNF143 is a widely expressed zinc-finger transcription factor that has been found to anchor chromatin interactions that connect distal regulatory elements with gene promoters[32].

Two databases (HaploReg, CIS-BP TF) evaluate ZNF143 as having low or no binding to the minor (reference) allele of rs2105898 and very high affinity to the major (alternate) allele of rs2105898:

**CIS-BP (log score)**
Reference (T) allele: 4.459
Alternate (G) allele: 13.273

**HaploReg (log score)**
Reference (T) allele: -0.4
Alternate (G) allele: 11.5

ZNF143 is a recently identified component of complexes that maintain topologically associated domains (TADs) in concert with CTCF and cohesin (SMC1, SMC3, RAD21, STAG1/2), both of which also have numerous ChIP-seq peaks overlapping rs2105898. Specifically, ZNF143 has been found to directly bind and regulate promoter interaction with distal enhancers, congruous with the observation of numerous RNA polymerase ChIP-seq peaks at rs2105898 but with nearest promoter being 14.5kb away (*HLA-DQA1*, downstream). Furthermore, as this region lies in the genomic neighborhood of many genes for which rs2105898 is a multi-tissue eQTL (*HLA-DRB1*, *-DRB5*, *-DRB6* upstream and *-DQA1*, *-DQA2*, *-DQB1*, and *-DQB2* downstream) it seems plausible that by regulating ZNF143 binding, rs2105898 alters the interaction between this enhancer region and the promoters of the numerous proximal *HLA* class II genes.

**rs2105898 is in strong LD with peak SNPs for other autoimmune disorders**

rs2105898 also has high LD to the most strongly associated SNPs for other autoimmune phenotypes. Of these associations, the strongest is to the peak SNP for multiple sclerosis oligoclonal band status ($r^2$=0.88,D'=0.98). Also in high LD to rs2105898 is a shared peak SNP for associations to broad multiple sclerosis, immunoglobulin A production, ulcerative colitis, and Crohn's disease (all $r^2$=0.49,D'=0.98).

**Schizophrenia Working Group of the Psychiatric Genomics Consortium**

Stephan Ripke[1,2], Benjamin M. Neale[1,2,3,4], Aiden Corvin[5], James T. R. Walters[6], Kai-How Farh[1], Peter A. Holmans[6,7], Phil Lee[1,2,4], Brendan Bulik-Sullivan[1,2], David A. Collier[8,9], Hailiang Huang[1,3], Tune H. Pers[3,10,11], Ingrid Agartz[12,13,14], Esben Agerbo[15,16,17], Margot Albus[18], Madeline Alexander[19], Farooq Amin[20,21], Silviu A. Bacanu[22], Martin Begemann[23], Richard A Belliveau Jr[2], Judit Bene[24,25], Sarah E. Bergen [2,26], Elizabeth Bevilacqua[2], Tim B Bigdeli [22], Donald W. Black[27], Richard Bruggeman[28], Nancy G. Buccola[29], Randy L. Buckner[30,31,32], William Byerley[33], Wiepke Cahn[34], Guiqing Cai[35,36], Murray J. Cairns[39,120,170], Dominique Campion[37], Rita M. Cantor[38], Vaughan J. Carr[39,40], Noa Carrera[6], Stanley V. Catts[39,41], Kimberly D. Chambert[2], Raymond C. K. Chan[42], Ronald Y. L. Chen[43], Eric Y. H. Chen[43,44], Wei Cheng[45], Eric F. C. Cheung[46], Siow Ann Chong[47], C. Robert Cloninger[48], David Cohen[49], Nadine Cohen[50], Paul Cormican[5], Nick Craddock[6,7], Benedicto Crespo-Facorro[210], James J. Crowley[51], David Curtis[52,53], Michael Davidson[54], Kenneth L. Davis[36], Franziska Degenhardt[55,56], Jurgen Del Favero[57], Lynn E. DeLisi[128,129] , Ditte Demontis[17,58,59], Dimitris Dikeos[60], Timothy Dinan[61], Srdjan Djurovic[14,62], Gary Donohoe[5,63], Elodie Drapeau[36], Jubao Duan[64,65], Frank Dudbridge[66], Naser Durmishi[67], Peter Eichhammer[68], Johan Eriksson[69,70,71], Valentina Escott-Price[6], Laurent Essioux[72], Ayman H. Fanous[73,74,75,76], Martilias S. Farrell[51], Josef Frank[77], Lude Franke[78], Robert Freedman[79], Nelson B. Freimer[80], Marion Friedl[81], Joseph I. Friedman[36], Menachem Fromer[1,2,4,82], Giulio Genovese[2], Lyudmila Georgieva[6], Elliot S. Gershon[209], Ina Giegling[81,83], Paola Giusti-Rodríguez[51], Stephanie Godard[84], Jacqueline I. Goldstein[1,3], Vera Golimbet[85], Srihari Gopal[86], Jacob Gratten[87], Lieuwe de Haan[88], Marina Mitjans[23], Marian L. Hamshere[6], Mark Hansen[89], Thomas Hansen[17,90], Vahram Haroutunian[36,91,92], Annette M. Hartmann[81], Frans A. Henskens[39,93,94], Stefan Herms[55,56,95], Joel N. Hirschhorn[3,11,96], Per Hoffmann[55,56,95], Andrea Hofman[55,56], Mads V. Hollegaard[97], David M. Hougaard[97], Masashi Ikeda[98], Inge Joa[99], Antonio Julià[100], René S. Kahn[34], Luba Kalaydjieva[101,102], Sena Karachanak-Yankova[103], Juha Karjalainen[78], David Kavanagh[6], Matthew C. Keller[104], Brian J. Kelly[120], James L. Kennedy[105,106,107], Andrey Khrunin[108], Yunjung Kim[51], Janis Klovins[109], James A. Knowles[110], Bettina Konte[81], Vaidutis Kucinskas[111], Zita Ausrele Kucinskiene[111], Hana Kuzelova-Ptackova[112], Anna K. Kähler[26], Claudine Laurent[19,113], Jimmy Lee Chee Keong[47,114], S. Hong Lee[87], Sophie E. Legge[6], Bernard Lerer[115], Miaoxin Li[43,44,116] Tao Li[117], Kung-Yee Liang[118], Jeffrey Lieberman[119], Svetlana Limborska[108], Carmel M. Loughland[39,120], Jan Lubinski[121], Jouko Lönnqvist[122], Milan Macek Jr[112], Patrik K. E. Magnusson[26], Brion S. Maher[123], Wolfgang Maier[124], Jacques Mallet[125], Sara Marsal[100], Manuel Mattheisen[17,58,59,126], Morten Mattingsdal[14,127], Robert W. McCarley[128,129], Colm McDonald[130], Andrew M. McIntosh[131,132], Sandra Meier[77], Carin J. Meijer[88], Bela Melegh[24,25], Ingrid Melle[14,133], Raquelle I. Mesholam-Gately[128,134], Andres Metspalu[135], Patricia T. Michie[39,136], Lili Milani[135], Vihra Milanova[137], Younes Mokrab[8], Derek W. Morris[5,63], Ole Mors[17,58,138], Kieran C. Murphy[139], Robin M. Murray[140], Inez Myin-Germeys[141], Bertram Müller-Myhsok[142,143,144], Mari Nelis[135], Igor Nenadic[145], Deborah A. Nertney[146], Gerald Nestadt[147], Kristin K. Nicodemus[148], Liene Nikitina-Zake[109], Laura Nisenbaum[149], Annelie Nordin[150], Eadbhard O'Callaghan[151], Colm O'Dushlaine[2], F. Anthony O'Neill[152], Sang-Yun Oh[153], Ann Olincy[79], Line Olsen[17,90], Jim Van Os[141,154], Psychosis Endophenotypes International Consortium[155], Christos Pantelis[39,156], George N. Papadimitriou[60], Agnes A. Steixner[23], Elena Parkhomenko[36], Michele T. Pato[110], Tiina Paunio[157,158], Milica Pejovic-Milovancevic[159], Diana O. Perkins[160], Olli Pietiläinen[158,161], Jonathan Pimm[53], Andrew J. Pocklington[6], John Powell[140], Alkes Price[3],[162], Ann E. Pulver[147], Shaun M. Purcell[82], Digby Quested[163], Henrik B. Rasmussen[17,90], Abraham Reichenberg[36], Mark A. Reimers[164], Alexander L. Richards[6], Joshua L. Roffman[30,32], Panos Roussos[82,165], Douglas M. Ruderfer[6,82], Veikko Salomaa[71], Alan R. Sanders[64,65], Ulrich Schall[39,120], Christian R. Schubert[166], Thomas G. Schulze[77,167], Sibylle G. Schwab[168], Edward M. Scolnick[2], Rodney J. Scott[39,169,170], Larry J. Seidman[128,134], Jianxin Shi[171], Engilbert Sigurdsson[172], Teimuraz Silagadze[173], Jeremy M. Silverman[36,174], Kang Sim[47], Petr Slominsky[108], Jordan W. Smoller[2,4], Hon-Cheong So[43], Chris C. A. Spencer[175], Eli A. Stahl[3,82], Hreinn Stefansson[176], Stacy Steinberg[176], Elisabeth Stogmann[177], Richard E. Straub[178], Eric Strengman[179,34], Jana Strohmaier[77], T. Scott Stroup[119], Mythily Subramaniam[47], Jaana Suvisaari[122], Dragan M. Svrakic[48], Jin P. Szatkiewicz[51], Erik Söderman[12], Srinivas Thirumalai[180], Draga Toncheva[103], Paul A. Tooney[39,120,170] , Sarah Tosato[181], Juha Veijola[182,183], John Waddington[184], Dermot Walsh[185], Dai Wang[86], Qiang Wang[117],

Bradley T. Webb[22], Mark Weiser[54], Dieter B. Wildenauer[186], Nigel M. Williams[6], Stephanie Williams[51], Stephanie H. Witt[77], Aaron R. Wolen[164], Emily H. M. Wong[43], Brandon K. Wormley[22], Jing Qin Wu[39,170], Hualin Simon Xi[187], Clement C. Zai[105,106], Xuebin Zheng[188], Fritz Zimprich[177], Naomi R. Wray[87], Kari Stefansson[176], Peter M. Visscher[87], Wellcome Trust Case-Control Consortium 2[189], Rolf Adolfsson[150], Ole A. Andreassen[14,133], Douglas H. R. Blackwood[132], Elvira Bramon[190], Joseph D. Buxbaum[35,36,91,191], Anders D. Børglum[17,58,59,138], Sven Cichon[55,56,95,192], Ariel Darvasi[193], Enrico Domenici[194], Hannelore Ehrenreich[23], Tõnu Esko[3,11,96,135], Pablo V. Gejman[64,65], Michael Gill[5], Hugh Gurling[53], Christina M. Hultman[26], Nakao Iwata[98], Assen V. Jablensky[39,102,186,195], Erik G. Jönsson[12,14], Kenneth S. Kendler[196], George Kirov[6], Jo Knight[105,106,107], Todd Lencz[197,198,199], Douglas F. Levinson[19], Qingqin S. Li[86], Jianjun Liu[188,200], Anil K. Malhotra[197,198,199], Steven A. McCarroll[2,96], Andrew McQuillin[53], Jennifer L. Moran[2], Preben B. Mortensen[15,16,17], Bryan J. Mowry[87,201], Markus M. Nöthen[55,56], Roel A. Ophoff[38,80,34], Michael J. Owen[6,7], Aarno Palotie[2,4,161], Carlos N. Pato[110], Tracey L. Petryshen[2,128,202], Danielle Posthuma[203,204,205], Marcella Rietschel[77], Brien P. Riley[196], Dan Rujescu[81,83], Pak C. Sham[43,44,116] Pamela Sklar[82,91,165], David St Clair[206], Daniel R. Weinberger[178,207], Jens R. Wendland[166], Thomas Werge[17,90,208], Mark J. Daly[1,2,3], Patrick F. Sullivan[26,51,160] & Michael C. O'Donovan[6,7]

[1]Analytic and Translational Genetics Unit, Massachusetts General Hospital, Boston, Massachusetts 02114, USA.

[2]Stanley Center for Psychiatric Research, Broad Institute of MIT and Harvard, Cambridge, Massachusetts 02142, USA.

[3]Medical and Population Genetics Program, Broad Institute of MIT and Harvard, Cambridge, Massachusetts 02142, USA.

[4]Psychiatric and Neurodevelopmental Genetics Unit, Massachusetts General Hospital, Boston, Massachusetts 02114, USA.

[5]Neuropsychiatric Genetics Research Group, Department of Psychiatry, Trinity College Dublin, Dublin 8, Ireland.

[6]MRC Centre for Neuropsychiatric Genetics and Genomics, Institute of Psychological Medicine and Clinical Neurosciences, School of Medicine, Cardiff University, Cardiff, CF24 4HQ, UK.

[7]National Centre for Mental Health, Cardiff University, Cardiff, CF24 4HQ, UK.

[8]Eli Lilly and Company Limited, Erl Wood Manor, Sunninghill Road, Windlesham, Surrey, GU20 6PH, UK. [9]Social, Genetic and Developmental Psychiatry Centre, Institute of Psychiatry, King's College London, London, SE5 8AF, UK.

[10]Center for Biological Sequence Analysis, Department of Systems Biology, Technical University of Denmark, DK-2800, Denmark.

[11]Division of Endocrinology and Center for Basic and Translational Obesity Research, Boston Children's Hospital, Boston, Massachusetts, 02115USA.

[12]Department of Clinical Neuroscience, Psychiatry Section, Karolinska Institutet, SE-17176 Stockholm, Sweden. [13]Department of Psychiatry, Diakonhjemmet Hospital, 0319 Oslo, Norway. [14]NORMENT, KG Jebsen Centre for Psychosis Research, Institute of Clinical Medicine, University of Oslo, 0424 Oslo, Norway.

[15]Centre for Integrative Register-based Research, CIRRAU, Aarhus University, DK-8210 Aarhus, Denmark.

[16]National Centre for Register-based Research, Aarhus University, DK-8210 Aarhus, Denmark.

[17]The Lundbeck Foundation Initiative for Integrative Psychiatric Research, iPSYCH, Denmark.

[18]State Mental Hospital, 85540 Haar, Germany.

[19]Department of Psychiatry and Behavioral Sciences, Stanford University, Stanford, California 94305, USA.

[20]Department of Psychiatry and Behavioral Sciences, Atlanta Veterans Affairs Medical Center, Atlanta, Georgia 30033, USA.

[21]Department of Psychiatry and Behavioral Sciences, Emory University, Atlanta Georgia 30322, USA.

[22]Virginia Institute for Psychiatric and Behavioral Genetics, Department of Psychiatry, Virginia Commonwealth University, Richmond, Virginia 23298, USA.

[23]Clinical Neuroscience, Max Planck Institute of Experimental Medicine, Göttingen 37075, Germany.

[24]Department of Medical Genetics, University of Pécs, Pécs H-7624, Hungary.

[25]Szentagothai Research Center, University of Pécs, Pécs H-7624, Hungary.

[26]Department of Medical Epidemiology and Biostatistics, Karolinska Institutet, Stockholm SE-17177, Sweden.

[27]Department of Psychiatry, University of Iowa Carver College of Medicine, Iowa City, Iowa 52242, USA.

[28]University Medical Center Groningen, Department of Psychiatry, University of Groningen NL-9700 RB, The Netherlands.

[29]School of Nursing, Louisiana State University Health Sciences Center, New Orleans, Louisiana 70112, USA.

[30]Athinoula A. Martinos Center, Massachusetts General Hospital, Boston, Massachusetts 02129, USA.

[31]Center for Brain Science, Harvard University, Cambridge, Massachusetts, 02138 USA.

[32]Department of Psychiatry, Massachusetts General Hospital, Boston, Massachusetts, 02114 USA.

[33]Department of Psychiatry, University of California at San Francisco, San Francisco, California, 94143 USA.

[34]University Medical Center Utrecht, Department of Psychiatry, Rudolf Magnus Institute of Neuroscience, 3584 Utrecht, The Netherlands.

[35]Department of Human Genetics, Icahn School of Medicine at Mount Sinai, New York, New York 10029 USA.

[36]Department of Psychiatry, Icahn School of Medicine at Mount Sinai, New York, New York 10029 USA.

[37]Centre Hospitalier du Rouvray and INSERM U1079 Faculty of Medicine, 76301 Rouen, France.

[38]Department of Human Genetics, David Geffen School of Medicine, University of California, Los Angeles, California 90095, USA.

[39]Schizophrenia Research Institute, Sydney NSW 2010, Australia.

[40]School of Psychiatry, University of New South Wales, Sydney NSW 2031, Australia.

[41]Royal Brisbane and Women’s Hospital, University of Queensland, Brisbane, St Lucia QLD 4072, Australia.

[42]Institute of Psychology, Chinese Academy of Science, Beijing 100101, China.

[43]Department of Psychiatry, Li Ka Shing Faculty of Medicine, The University of Hong Kong, Hong Kong, China.

[44]State Key Laboratory for Brain and Cognitive Sciences, Li Ka Shing Faculty of Medicine, The University of Hong Kong, Hong Kong, China.

[45]Department of Computer Science, University of North Carolina, Chapel Hill, North Carolina 27514, USA.

[46]Castle Peak Hospital, Hong Kong, China.

[47]Institute of Mental Health, Singapore 539747, Singapore.

[48]Department of Psychiatry, Washington University, St. Louis, Missouri 63110, USA.

[49]Department of Child and Adolescent Psychiatry, Assistance Publique Hopitaux de Paris, Pierre and Marie Curie Faculty of Medicine and Institute for Intelligent Systems and Robotics, Paris, 75013, France.

[50] Blue Note Biosciences, Princeton, New Jersey 08540, USA

[51]Department of Genetics, University of North Carolina, Chapel Hill, North Carolina 27599-7264, USA.

[52]Department of Psychological Medicine, Queen Mary University of London, London E1 1BB, UK.

[53]Molecular Psychiatry Laboratory, Division of Psychiatry, University College London, London WC1E 6JJ, UK.

[54]Sheba Medical Center, Tel Hashomer 52621, Israel.

[55]Department of Genomics, Life and Brain Center, D-53127 Bonn, Germany.

[56]Institute of Human Genetics, University of Bonn, D-53127 Bonn, Germany.

[57]Applied Molecular Genomics Unit, VIB Department of Molecular Genetics, University of Antwerp, B-2610 Antwerp, Belgium.

[58]Centre for Integrative Sequencing, iSEQ, Aarhus University, DK-8000 Aarhus C, Denmark.

[59]Department of Biomedicine, Aarhus University, DK-8000 Aarhus C, Denmark.

[60]First Department of Psychiatry, University of Athens Medical School, Athens 11528, Greece.

[61]Department of Psychiatry, University College Cork, Co. Cork, Ireland.

[62]Department of Medical Genetics, Oslo University Hospital, 0424 Oslo, Norway.

[63]Cognitive Genetics and Therapy Group, School of Psychology and Discipline of Biochemistry, National University of Ireland Galway, Co. Galway, Ireland.

[64]Department of Psychiatry and Behavioral Neuroscience, University of Chicago, Chicago, Illinois 60637, USA.

[65]Department of Psychiatry and Behavioral Sciences, NorthShore University HealthSystem, Evanston, Illinois 60201, USA.

[66]Department of Non-Communicable Disease Epidemiology, London School of Hygiene and Tropical Medicine, London WC1E 7HT, UK.

[67]Department of Child and Adolescent Psychiatry, University Clinic of Psychiatry, Skopje 1000, Republic of Macedonia.

[68]Department of Psychiatry, University of Regensburg, 93053 Regensburg, Germany.

[69]Department of General Practice, Helsinki University Central Hospital, University of Helsinki P.O. Box 20, Tukholmankatu 8 B, FI-00014, Helsinki, Finland

[70]Folkhälsan Research Center, Helsinki, Finland, Biomedicum Helsinki 1, Haartmaninkatu 8, FI-00290, Helsinki, Finland.

[71]National Institute for Health and Welfare, P.O. BOX 30, FI-00271 Helsinki, Finland.

[72]Translational Technologies and Bioinformatics, Pharma Research and Early Development, F. Hoffman-La Roche, CH-4070 Basel, Switzerland.

[73]Department of Psychiatry, Georgetown University School of Medicine, Washington DC 20057, USA.

[74]Department of Psychiatry, Keck School of Medicine of the University of Southern California, Los Angeles, California 90033, USA.

[75]Department of Psychiatry, Virginia Commonwealth University School of Medicine, Richmond, Virginia 23298, USA.

[76]Mental Health Service Line, Washington VA Medical Center, Washington DC 20422, USA.

[77]Department of Genetic Epidemiology in Psychiatry, Central Institute of Mental Health, Medical Faculty Mannheim, University of Heidelberg, Heidelberg , D-68159 Mannheim, Germany.

[78]Department of Genetics, University of Groningen, University Medical Centre Groningen, 9700 RB Groningen, The Netherlands.

[79]Department of Psychiatry, University of Colorado Denver, Aurora, Colorado 80045, USA.

[80]Center for Neurobehavioral Genetics, Semel Institute for Neuroscience and Human Behavior, University of California, Los Angeles, California 90095, USA.

[81]Department of Psychiatry, University of Halle, 06112 Halle, Germany.

[82]Division of Psychiatric Genomics, Department of Psychiatry, Icahn School of Medicine at Mount Sinai, New York, New York 10029, USA.

[83]Department of Psychiatry, University of Munich, 80336, Munich, Germany.

[84]Departments of Psychiatry and Human and Molecular Genetics, INSERM, Institut de Myologie, Hôpital de la Pitiè-Salpêtrière, Paris, 75013, France.

[85]Mental Health Research Centre, Russian Academy of Medical Sciences, 115522 Moscow, Russia.

[86]Neuroscience Therapeutic Area, Janssen Research and Development, Raritan, New Jersey 08869, USA.

[87]Queensland Brain Institute, The University of Queensland, Brisbane, Queensland, QLD 4072, Australia.

[88]Academic Medical Centre University of Amsterdam, Department of Psychiatry, 1105 AZ Amsterdam, The Netherlands.

[89]Illumina, La Jolla, California, California 92122, USA.

[90]Institute of Biological Psychiatry, Mental Health Centre Sct. Hans, Mental Health Services Copenhagen, DK-4000, Denmark.

[91]Friedman Brain Institute, Icahn School of Medicine at Mount Sinai, New York, New York 10029, USA.

[92]J. J. Peters VA Medical Center, Bronx, New York, New York 10468, USA.

[93]Priority Research Centre for Health Behaviour, University of Newcastle, Newcastle NSW 2308, Australia.

[94]School of Electrical Engineering and Computer Science, University of Newcastle, Newcastle NSW 2308, Australia.

[95]Division of Medical Genetics, Department of Biomedicine, University of Basel, Basel, CH-4058, Switzerland.

[96]Department of Genetics, Harvard Medical School, Boston, Massachusetts 02115, USA.

[97]Section of Neonatal Screening and Hormones, Department of Clinical Biochemistry, Immunology and Genetics, Statens Serum Institut, Copenhagen, DK-2300, Denmark.

[98]Department of Psychiatry, Fujita Health University School of Medicine, Toyoake, Aichi, 470-1192, Japan.

[99]Regional Centre for Clinical Research in Psychosis, Department of Psychiatry, Stavanger University Hospital, 4011 Stavanger, Norway.

[100]Rheumatology Research Group, Vall d'Hebron Research Institute, Barcelona, 08035, Spain.

[101]Centre for Medical Research, The University of Western Australia, Perth, WA 6009, Australia.

[102]The Perkins Institute for Medical Research, The University of Western Australia, Perth, WA 6009, Australia.

[103]Department of Medical Genetics, Medical University, Sofia1431, Bulgaria.

[104]Department of Psychology, University of Colorado Boulder, Boulder, Colorado 80309, USA.

[105]Campbell Family Mental Health Research Institute, Centre for Addiction and Mental Health, Toronto, Ontario, M5T 1R8, Canada.

[106]Department of Psychiatry, University of Toronto, Toronto, Ontario, M5T 1R8, Canada.

[107]Institute of Medical Science, University of Toronto, Toronto, Ontario, M5S 1A8, Canada.

[108]Institute of Molecular Genetics, Russian Academy of Sciences, Moscow123182, Russia.

[109]Latvian Biomedical Research and Study Centre, Riga, LV-1067, Latvia.

[110]Department of Psychiatry and Zilkha Neurogenetics Institute, Keck School of Medicine at University of Southern California, Los Angeles, California 90089, USA.

[111]Faculty of Medicine, Vilnius University, LT-01513 Vilnius, Lithuania.

[112] Department of Biology and Medical Genetics, 2nd Faculty of Medicine and University Hospital Motol, 150 06 Prague, Czech Republic.

[113] Department of Child and Adolescent Psychiatry, Pierre and Marie Curie Faculty of Medicine, Paris 75013, France.

[114]Duke-NUS Graduate Medical School, Singapore 169857, Singapore.

[115]Department of Psychiatry, Hadassah-Hebrew University Medical Center, Jerusalem 91120, Israel.

[116]Centre for Genomic Sciences, The University of Hong Kong, Hong Kong, China.

[117]Mental Health Centre and Psychiatric Laboratory, West China Hospital, Sichuan University, Chengdu, 610041, Sichuan, China.

[118]Department of Biostatistics, Johns Hopkins University Bloomberg School of Public Health, Baltimore, Maryland 21205, USA.

[119]Department of Psychiatry, Columbia University, New York, New York 10032, USA.

[120]Priority Centre for Translational Neuroscience and Mental Health, University of Newcastle, Newcastle NSW 2300, Australia.

[121]Department of Genetics and Pathology, International Hereditary Cancer Center, Pomeranian Medical University in Szczecin, 70-453 Szczecin, Poland.

[122]Department of Mental Health and Substance Abuse Services; National Institute for Health and Welfare, P.O. BOX 30, FI-00271 Helsinki, Finland

[123]Department of Mental Health, Bloomberg School of Public Health, Johns Hopkins University, Baltimore, Maryland 21205, USA.

[124]Department of Psychiatry, University of Bonn, D-53127 Bonn, Germany.

[125]Centre National de la Recherche Scientifique, Laboratoire de Génétique Moléculaire de la Neurotransmission et des Processus Neurodégénératifs, Hôpital de la Pitié Salpêtrière, 75013, Paris, France.

[126]Department of Genomics Mathematics, University of Bonn, D-53127 Bonn, Germany.

[127]Research Unit, Sørlandet Hospital, 4604 Kristiansand, Norway.

[128]Department of Psychiatry, Harvard Medical School, Boston, Massachusetts 02115, USA.

[129]VA Boston Health Care System, Brockton, Massachusetts 02301, USA.

[130]Department of Psychiatry, National University of Ireland Galway, Co. Galway, Ireland.

[131]Centre for Cognitive Ageing and Cognitive Epidemiology, University of Edinburgh, Edinburgh EH16 4SB, UK.

[132]Division of Psychiatry, University of Edinburgh, Edinburgh EH16 4SB, UK.

[133]Division of Mental Health and Addiction, Oslo University Hospital, 0424 Oslo, Norway.

[134]Massachusetts Mental Health Center Public Psychiatry Division of the Beth Israel Deaconess Medical Center, Boston, Massachusetts 02114, USA.

[135]Estonian Genome Center, University of Tartu, Tartu 50090, Estonia.

[136]School of Psychology, University of Newcastle, Newcastle NSW 2308, Australia.

[137]First Psychiatric Clinic, Medical University, Sofia 1431, Bulgaria.

[138]Department P, Aarhus University Hospital, DK-8240 Risskov, Denmark.

[139]Department of Psychiatry, Royal College of Surgeons in Ireland, Dublin 2, Ireland.

[140]King’s College London, London SE5 8AF, UK.

[141]Maastricht University Medical Centre, South Limburg Mental Health Research and Teaching Network, EURON, 6229 HX Maastricht, The Netherlands.

[142]Institute of Translational Medicine, University of Liverpool, Liverpool L69 3BX, UK.

[143]Max Planck Institute of Psychiatry, 80336 Munich, Germany.

[144]Munich Cluster for Systems Neurology (SyNergy), 80336 Munich, Germany.

[145]Department of Psychiatry and Psychotherapy, Jena University Hospital, 07743 Jena, Germany.

[146]Department of Psychiatry, Queensland Brain Institute and Queensland Centre for Mental Health Research, University of Queensland, Brisbane, Queensland, St Lucia QLD 4072, Australia.

[147]Department of Psychiatry and Behavioral Sciences, Johns Hopkins University School of Medicine, Baltimore, Maryland 21205, USA.

[148]Department of Psychiatry, Trinity College Dublin, Dublin 2, Ireland.

[149]Eli Lilly and Company, Lilly Corporate Center, Indianapolis, 46285 Indiana, USA.

[150]Department of Clinical Sciences, Psychiatry, Umeå University, SE-901 87 Umeå, Sweden.

[151]DETECT Early Intervention Service for Psychosis, Blackrock, Co. Dublin, Ireland.

[152]Centre for Public Health, Institute of Clinical Sciences, Queen's University Belfast, Belfast BT12 6AB, UK.

[153]Lawrence Berkeley National Laboratory, University of California at Berkeley, Berkeley, California 94720, USA.

[154]Institute of Psychiatry, King's College London, London SE5 8AF, UK.

[155]A list of authors and affiliations appear in the Supplementary Information.

[156]Melbourne Neuropsychiatry Centre, University of Melbourne & Melbourne Health, Melbourne, Vic 3053, Australia.

[157]Department of Psychiatry, University of Helsinki, P.O. Box 590, FI-00029 HUS, Helsinki, Finland.

[158]Public Health Genomics Unit, National Institute for Health and Welfare, P.O. BOX 30, FI-00271 Helsinki, Finland.

[159]Medical Faculty, University of Belgrade, 11000 Belgrade, Serbia.

[160]Department of Psychiatry, University of North Carolina, Chapel Hill, North Carolina 27599-7160, USA.

[161]Institute for Molecular Medicine Finland, FIMM, University of Helsinki, P.O. Box 20

FI-00014, Helsinki, Finland.

[162]Department of Epidemiology, Harvard School of Public Health, Boston, Massachusetts 02115, USA.

[163]Department of Psychiatry, University of Oxford, Oxford, OX3 7JX, UK.

[164]Virginia Institute for Psychiatric and Behavioral Genetics, Virginia Commonwealth University, Richmond, Virginia 23298, USA.

[165]Institute for Multiscale Biology, Icahn School of Medicine at Mount Sinai, New York, New York 10029, USA.

[166]PharmaTherapeutics Clinical Research, Pfizer Worldwide Research and Development, Cambridge, Massachusetts 02139, USA.

[167]Department of Psychiatry and Psychotherapy, University of Gottingen, 37073 Göttingen, Germany.

[168]Psychiatry and Psychotherapy Clinic, University of Erlangen, 91054 Erlangen, Germany.

[169]Hunter New England Health Service, Newcastle NSW 2308, Australia.

[170]School of Biomedical Sciences and Pharmacy, University of Newcastle, Callaghan NSW 2308, Australia.

[171]Division of Cancer Epidemiology and Genetics, National Cancer Institute, Bethesda, Maryland 20892, USA.

[172]University of Iceland, Landspitali, National University Hospital, 101 Reykjavik, Iceland.

[173]Department of Psychiatry and Drug Addiction, Tbilisi State Medical University (TSMU), **N33, 0177** Tbilisi, Georgia.

[174]Research and Development, Bronx Veterans Affairs Medical Center, New York, New York 10468, USA.

[175]Wellcome Trust Centre for Human Genetics, Oxford, OX3 7BN, UK.

[176]deCODE Genetics, 101 Reykjavik, Iceland.

[177]Department of Clinical Neurology, Medical University of Vienna, 1090 Wien, Austria.

[178]Lieber Institute for Brain Development, Baltimore, Maryland 21205, USA.

[179]Department of Medical Genetics, University Medical Centre Utrecht, Universiteitsweg 100, 3584 CG, Utrecht, The Netherlands.

[180]Berkshire Healthcare NHS Foundation Trust, Bracknell RG12 1BQ, UK.

[181]Section of Psychiatry, University of Verona, 37134 Verona, Italy.

[182]Department of Psychiatry, University of Oulu, P.O. BOX 5000, 90014, Finland

[183]University Hospital of Oulu, P.O.BOX 20, 90029 OYS, Finland.

[184]Molecular and Cellular Therapeutics, Royal College of Surgeons in Ireland, Dublin 2, Ireland.

[185]Health Research Board, Dublin 2, Ireland.

[186]School of Psychiatry and Clinical Neurosciences, The University of Western Australia, Perth WA6009, Australia.

[187]Computational Sciences CoE, Pfizer Worldwide Research and Development, Cambridge, Massachusetts 02139, USA.

[188]Human Genetics, Genome Institute of Singapore, A*STAR, Singapore 138672, Singapore.

[190]University College London, London WC1E 6BT, UK.

[191]Department of Neuroscience, Icahn School of Medicine at Mount Sinai, New York, New York 10029, USA.

[192]Institute of Neuroscience and Medicine (INM-1), Research Center Juelich, 52428 Juelich, Germany.

[193]Department of Genetics, The Hebrew University of Jerusalem, 91905 Jerusalem, Israel.

[194]Neuroscience Discovery and Translational Area, Pharma Research and Early Development, F. Hoffman-La Roche, CH-4070 Basel, Switzerland.

[195]Centre for Clinical Research in Neuropsychiatry, School of Psychiatry and Clinical Neurosciences, The University of Western Australia, Medical Research Foundation Building, Perth WA 6000, Australia.

[196]Virginia Institute for Psychiatric and Behavioral Genetics, Departments of Psychiatry and Human and Molecular Genetics, Virginia Commonwealth University, Richmond, Virginia 23298, USA.

[197]The Feinstein Institute for Medical Research, Manhasset, New York, 11030 USA.

[198]The Hofstra NS-LIJ School of Medicine, Hempstead, New York, 11549 USA.

[199]The Zucker Hillside Hospital, Glen Oaks, New York,11004 USA.

[200]Saw Swee Hock School of Public Health, National University of Singapore, Singapore 117597, Singapore.

[201]Queensland Centre for Mental Health Research, University of Queensland, Brisbane 4076, Queensland, Australia.

[202]Center for Human Genetic Research and Department of Psychiatry, Massachusetts General Hospital, Boston, Massachusetts 02114, USA.

[203]Department of Child and Adolescent Psychiatry, Erasmus University Medical Centre, Rotterdam 3000, The Netherlands.

[204]Department of Complex Trait Genetics, Neuroscience Campus Amsterdam, VU University Medical Center Amsterdam, Amsterdam 1081, The Netherlands.

[205]Department of Functional Genomics, Center for Neurogenomics and Cognitive Research, Neuroscience Campus Amsterdam, VU University, Amsterdam 1081, The Netherlands.

[206]University of Aberdeen, Institute of Medical Sciences, Aberdeen, AB25 2ZD, UK.

[207]Departments of Psychiatry, Neurology, Neuroscience and Institute of Genetic Medicine, Johns Hopkins School of Medicine, Baltimore, Maryland 21205, USA.

[208]Department of Clinical Medicine, University of Copenhagen, Copenhagen 2200, Denmark.

[209]Departments of Psychiatry and Human Genetics, University of Chicago, Chicago, Illinois 60637, USA.

[210]University Hospital Marqués de Valdecilla, Instituto de Formación e Investigación Marqués de Valdecilla, University of Cantabria, ED39008 Santander, Spain.
